# Asymmetric Hapln1a drives regionalised cardiac ECM expansion and promotes heart morphogenesis during zebrafish development

**DOI:** 10.1101/838128

**Authors:** Christopher J Derrick, Juliana Sánchez-Posada, Farah Hussein, Federico Tessadori, Eric JG Pollitt, Aaron M Savage, Robert N Wilkinson, Timothy J Chico, Fredericus J van Eeden, Jeroen Bakkers, Emily S Noël

**Affiliations:** Department of Biomedical Science, University of Sheffield, Western Bank, Sheffield, UK; Department of Infection, Immunity and Cardiovascular Disease, University of Sheffield, Western Bank, Sheffield, UK; School of Life Sciences, University of Nottingham, Queen’s Medical Centre, Nottingham, UK; Hubrecht Institute for Developmental and Stem Cell Biology, Uppsalalaan 8, Utrecht, NL

## Abstract

The mature vertebrate heart develops from a simple linear cardiac tube during early development through a series of highly asymmetric morphogenetic processes including cardiac looping and chamber ballooning. While the directionality of heart morphogenesis is partly controlled by embryonic laterality signals, previous studies have suggested that these extrinsic laterality cues interact with tissue-intrinsic signals in the heart to ensure robust asymmetric cardiac morphogenesis. Using live *in vivo* imaging of zebrafish embryos we describe a left-sided, chamber-specific expansion of the extracellular matrix (ECM) between the myocardium and endocardium at early stages of heart morphogenesis. We use Tomo-seq, a spatial transcriptomic approach, to identify transient and regionalised expression of *hyaluronan and proteoglycan link protein 1a* (*hapln1a),* encoding an ECM cross-linking protein, in the heart tube prior to cardiac looping overlapping with regionalised ECM expansion. Loss- and gain-of-function experiments demonstrate that regionalised Hapln1a promotes heart morphogenesis through regional modulation of ECM thickness in the heart tube. Finally, we show that while induction of asymmetric *hapln1a* expression is independent of embryonic left-right asymmetry, these laterality cues are required to orient the *hapln1a*-expressing cells asymmetrically along the left-right axis of the heart tube.

Together, we propose a model whereby laterality cues position *hapln1a* expression on the left of the heart tube, and this asymmetric Hapln1a deposition drives ECM asymmetry and subsequently promotes robust asymmetric cardiac morphogenesis.

## Introduction

Congenital heart defects are the most common human birth abnormality, with an incidence of approximately 1% of live births (van der Linde et al. 2011; Hoffman & Kaplan 2002). These structural malformations arise due to abnormal morphogenesis and maturation of the heart during embryonic development. A key stage in cardiac development is when the heart transitions from a linear tube to an asymmetric organ, a process including initial looping morphogenesis of the tube and subsequent ballooning of the cardiac chambers. Correct cardiac morphogenesis is vital for ensuring normal blood flow through the heart, proper chamber and vessel alignment, valve formation and septation. Therefore early cardiac morphogenesis is a tightly controlled process, and requires coordination of heart-extrinsic signalling cues, cardiac growth and tissue-intrinsic changes in cell shape, mediated by cytoskeletal rearrangements (Desgrange et al. 2018).

The requirement for embryonic left-right signalling pathways in promoting directionality of heart morphogenesis is well established, with asymmetric Nodal signalling playing a key role in driving rightward looping of the linear heart tube in multiple organisms (Levin et al. 1995; Lowe et al. 1996; Long 2003; Brennan et al. 2002; Toyoizumi et al. 2005). However, while embryos with defective asymmetric Nodal signalling display disrupted directionality of heart looping, the heart still undergoes looping morphogenesis (Noël et al. 2013; Brennan et al. 2002). This indicates that while extrinsic asymmetric cues provide directional information to the heart, regionalised intrinsic signals help to promote asymmetric morphogenesis. Supporting the model of tissue intrinsic heart looping, chick and zebrafish studies have demonstrated that heart tubes cultured *ex vivo* retain the ability to undergo aspects of morphogenesis without exposure to regionalised signalling or physical constraints from the embryo (Noël et al. 2013; Manning & McLachlan 1990). It is likely that the interplay of extrinsic and intrinsic regionalised signaling and cell behaviours ensure the coordination of directionality and morphogenesis required to shape the heart and orient it within the body (Desgrange et al. 2018). The developing heart tube is composed of two cell layers: an outer tube of myocardium surrounding an inner layer of specialised endothelial cells (endocardium). Separating these two layers is the cardiac jelly, a specialised extracellular matrix (ECM). Cardiac jelly consists of collagens, glycosaminoglycans and glycoproteins and plays a pivotal role in providing mechanical cues and modulating extracellular signalling in the heart during cardiac development (Rozario & DeSimone 2010; Daley & Yamada 2013). Classic embryological experiments initially demonstrated that the cardiac jelly is important for heart morphogenesis (Barry, 1948; Nakamura and Manasek, 1981), while more recent studies have begun to identify specific ECM constituents with distinct roles in heart development (Rotstein et al. 2018; Chowdhury et al. 2017; Rambeau et al. 2017; Patra et al. 2011; Strate et al. 2015; Mittal et al. 2010; Tao et al. 2012; Wirrig et al. 2007; Trinh & Stainier 2004; Tsuda et al. 1998). Hyaluronic acid (HA) is a glycosaminoglycan with conserved roles in heart tube formation, cardiac morphogenesis and atrioventricular valve development (Camenisch et al. 2000; Smith et al. 2008; Chowdhury et al. 2017), suggesting a broad requirement for HA at various stages during cardiac development. However, the mechanisms by which the HA-rich ECM specifically promotes heart morphogenesis is unclear.

Recent studies in chick and mouse have reported that asymmetric ECM expansion in the mesoderm surrounding the gut tube promotes directional looping morphogenesis of the intestine (Sivakumar et al. 2018). This ECM expansion occurs through asymmetric modification of HA, raising interesting questions regarding the role of regional ECM remodeling and asymmetry in the morphogenesis of other tubular structures, including the heart.

In this study we demonstrate that the cardiac ECM of the zebrafish heart tube exhibits regionalised expansion prior to onset of looping morphogenesis, with an expanded ECM observed in both the left side and future atrium of the heart tube. Loss-of-function analyses demonstrate that this regionalised cardiac ECM expansion is dependent upon the ECM cross-linking protein Hyaluronan and Proteoglycan Link Protein 1a (Hapln1a), and that Hapln1a promotes heart morphogenesis. Finally, we show that while asymmetric *hapln1a* expression is independent of laterality cues, the axis of *hapln1a* asymmetry in the heart is dictated by embryonic laterality, and suggest a model where embryonic left-right asymmetry tightly defines the orientation of ECM asymmetry in the heart tube, and together these pathways promote asymmetric morphogenesis.

## Results

### The cardiac ECM is asymmetrically expanded at early stages of heart looping morphogenesis

During cardiac development the myocardial and endocardial layers of the heart are separated by a specialised ECM, the cardiac jelly. We hypothesised that there may be regional differences in the ECM of the zebrafish heart tube which drive looping morphogenesis. To examine regional ECM thickness in the heart tube we used live *in vivo* light-sheet microscopy to image quadruple transgenic zebrafish embryos at 26 hours post fertilisation (hpf).

*Tg(myl7:lifeActGFP); Tg(fli1a:AC-TagRFP); Tg(lft2BAC:Gal4FF); Tg(UAS:RFP)* zebrafish express actin-tagged GFP in the myocardium (Reischauer et al. 2014) and actin-localised RFP in the endothelium including the endocardium (Savage et al. 2019), allowing visualisation of the two tissue layers in the heart tube. The *Tg(lft2BAC:Gal4FF); Tg(UAS:RFP)* double transgenic drives RFP in *lefty2*-expressing cells, which comprises the dorsal myocardium of the heart tube at 26hpf (Fig 1A and Fig S1) (Smith et al. 2008; Baker et al. 2008). This combination of transgenes allowed imaging of optical cross sections through the heart tube at 26hpf, just prior to onset of looping morphogenesis, and enabled dorsal-ventral axis orientation of the heart tube (Fig 1A-G, Supplemental Movie 1). We consistently observed an asymmetry in the extracellular space between the myocardial and endocardial layers of the heart in the atrium, with an apparent thickening of the ECM on the left side of the tube which is maintained throughout the cardiac cycle (Fig 1D, G). With this combination of transgenes, we could quantify the extracellular space between the two tissue layers of the heart tube and calculate the ECM asymmetry ratio (left ECM thickness divided by right ECM thickness, where a value of >1 indicates a left sided expansion). Using this method, we detected a reproducible expansion of the ECM in the left side of the heart tube (Fig 1H). Further characterisation also found that the left-sided ECM expansion is maintained in the atrium at 50hpf (Fig S2).

**Figure 1.**
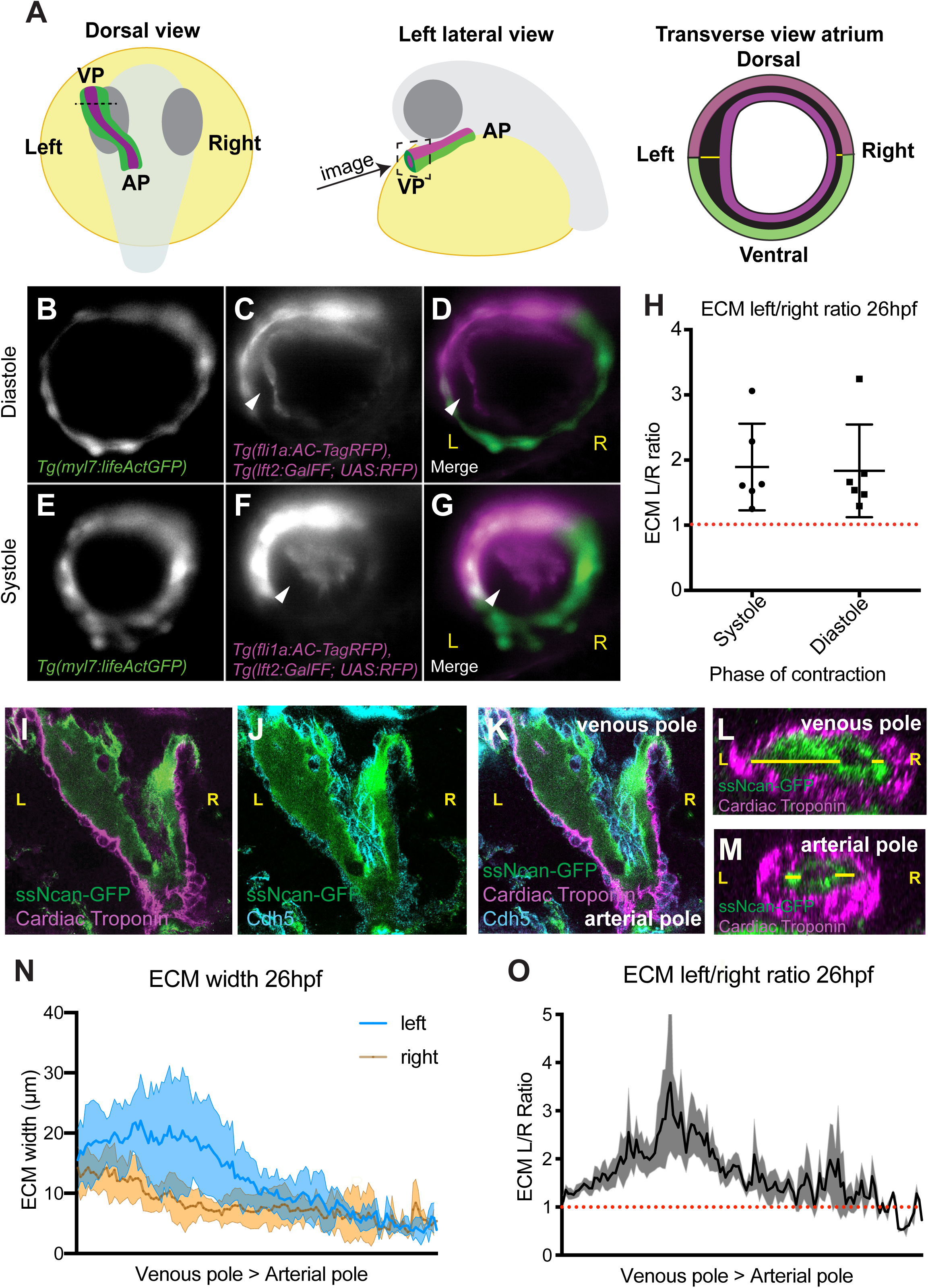
The hyaluronan-rich ECM is asymmetric during early heart development. A: Schematic depicting the developmental stage and orientation of embryos used in live imaging experiments. Optical transverse sections of the heart tube are imaged at the position of the dotted line/dotted square. Green - myocardium, magenta – endocardium, pink – dorsal myocardium. VP – venous pole, AP – arterial pole. B-G: Light sheet optical cross-sections through the heart tube of a 26hpf *Tg(myl7:lifeActGFP); Tg(fli1a:AC-TagRFP); Tg(lft2BAC:Gal4FF,UAS:RFP)* transgenic embryo during diastole (B-D) and systole (E-G) at the level of the dotted line in A. The myocardium is marked in green (B, D, E, G), and the dorsal myocardium and endocardium are marked in magenta (C, D, F, G). The extracellular space between the myocardium and endocardium is expanded on the left side of the heart tube (white arrowhead). H: Quantification of left-right ECM ratio in heart tubes, where a value greater than 1 (red dotted line) denotes left-sided expansion, n=6. I-K: Single confocal z-planes longitudinally through the heart at 26hpf of embryos injected with *ssNcan-GFP* (green), counterstained with cardiac troponin (magenta, I, K) and VE-Cadherin (cyan J, K). L-M: Transverse optical reslice through the 26hpf heart tube at the level of the venous pole (L) or arterial pole (M), ssNcan-GFP in green, cardiac troponin in magenta. ECM width is measured using the ssNcan-GFP signal (yellow line) on left and right sides of the tube. N: Quantification of ECM width on the left (blue) and right (orange) sides of the heart tube from venous pole to arterial pole at 26hpf. Mean +/- SD are plotted, n=7. O: left-right ECM ratio in the heart tube from venous pole to arterial pole, where a value >1 (red dotted line) indicates a left-sided expansion. Mean +/- SD are plotted. L – left, R – right, VP – venous pole, AP – arterial pole.

Due to limited ability to image deeper cardiac tissue with sufficient resolution at 26hpf, we could only image the relatively superficially located venous pole/atrium of the heart tube in live embryos. Therefore, to determine whether ECM left-right asymmetry is restricted to the atrium of the heart tube or is maintained along the atrioventricular axis of the heart, we performed fixed tissue imaging. Previous studies have demonstrated that hyaluronic acid (HA) is present in the cardiac jelly during zebrafish heart development (Grassini et al. 2018; Camenisch et al. 2000; Lagendijk et al. 2011). To visualise the HA-rich ECM, wild type embryos were injected with the HA sensor *ssNcan-GFP* (De Angelis et al. 2017) at the 1-cell stage, fixed at 26hpf, and the GFP signal detected by immunohistochemistry before imaging the entire heart tube as a z-stack using confocal microscopy (Fig 1I-K). Optical reslicing of z-stacks generated cross sections of the heart tube from the venous pole to the arterial pole, allowing us to quantify the width of the *ssNcan-GFP*-positive ECM in the heart tube on both left and right sides of the tube along the entire pole-to-pole length of the heart (Fig 1L-N). We confirmed that the ECM is thicker on the left side of the heart tube compared the right, however this left/right asymmetry appears to be more profound in the venous pole/future atrium than in the arterial pole/future ventricle (Fig 1N). By employing the same quantitative analysis of ECM ratio as used in live imaging experiments, we observed that the ECM indeed exhibits the highest level of asymmetry in the future atrial region of the heart tube (Fig 1O). Together these data demonstrate that the heart tube exhibits an asymmetrically expanded ECM prior to onset of looping morphogenesis.

### hapln1a exhibits regionalised cardiac expression prior to heart morphogenesis

The asymmetric expansion of the cardiac ECM could be due to regionalised synthesis of ECM components. However, we did not observe any clear asymmetry in levels of HA deposition in the cardiac ECM in either live embryos injected with the *ssNcan-GFP* sensor (Fig S3) or in fixed hearts (Fig 1I-K). We also did not find any anteroposterior asymmetry in the heart disc, or left-right asymmetry in the heart tube in the expression of *hyaluronan synthase 2 (has2,* the major HA producing enzyme), *chondroitin sulfate synthase 1* (*chsy1*) or the ECM proteoglycans *versican* (*vcana/b), aggrecan (acana/b)*, all of which have previously been implicated in heart development (Peal et al. 2009; Camenisch et al. 2000; Smith et al. 2008; Rambeau et al. 2017; Mittal et al. 2019; Mjaatvedt et al. 1998) (Fig S3), suggesting that regionalised synthesis of these proteins does not cause ECM asymmetry. We therefore hypothesised that a protein required either for HA modification or cross-linking may be regionally expressed in the heart tube prior to looping morphogenesis and promote regionalised ECM expansion.

To identify candidate genes which modulate cardiac ECM expansion, we took a genome-wide approach to identify genes expressed in the heart tube at 26hpf, prior to the onset of looping morphogenesis. Since we observed the strongest left-sided ECM expansion in the putative atrium, as well as a generally more expanded ECM at the venous pole of the heart compared to the arterial pole, we used the previously described Tomo-seq technique to generate a regionalised map of gene expression from pole-to-pole in the heart tube (Junker et al. 2014; Burkhard & Bakkers 2018) (Figure 2A). We sectioned two individual hearts along the atrioventricular axis, identifying 6,787 and 8,916 expressed genes (see Supplementary Tables 2-5), of which approximately half were expressed in more than one section. By identifying which sections express the atrial marker *myh6* (*myosin, heavy chain 6, cardiac muscle, alpha*) we defined a subset of tissue sections with atrial identity. We subsequently filtered genes that are up-regulated in atrial sections compared to ventricular sections in both hearts and examined this list for genes which may be implicated in ECM modification.

**Figure 2.**
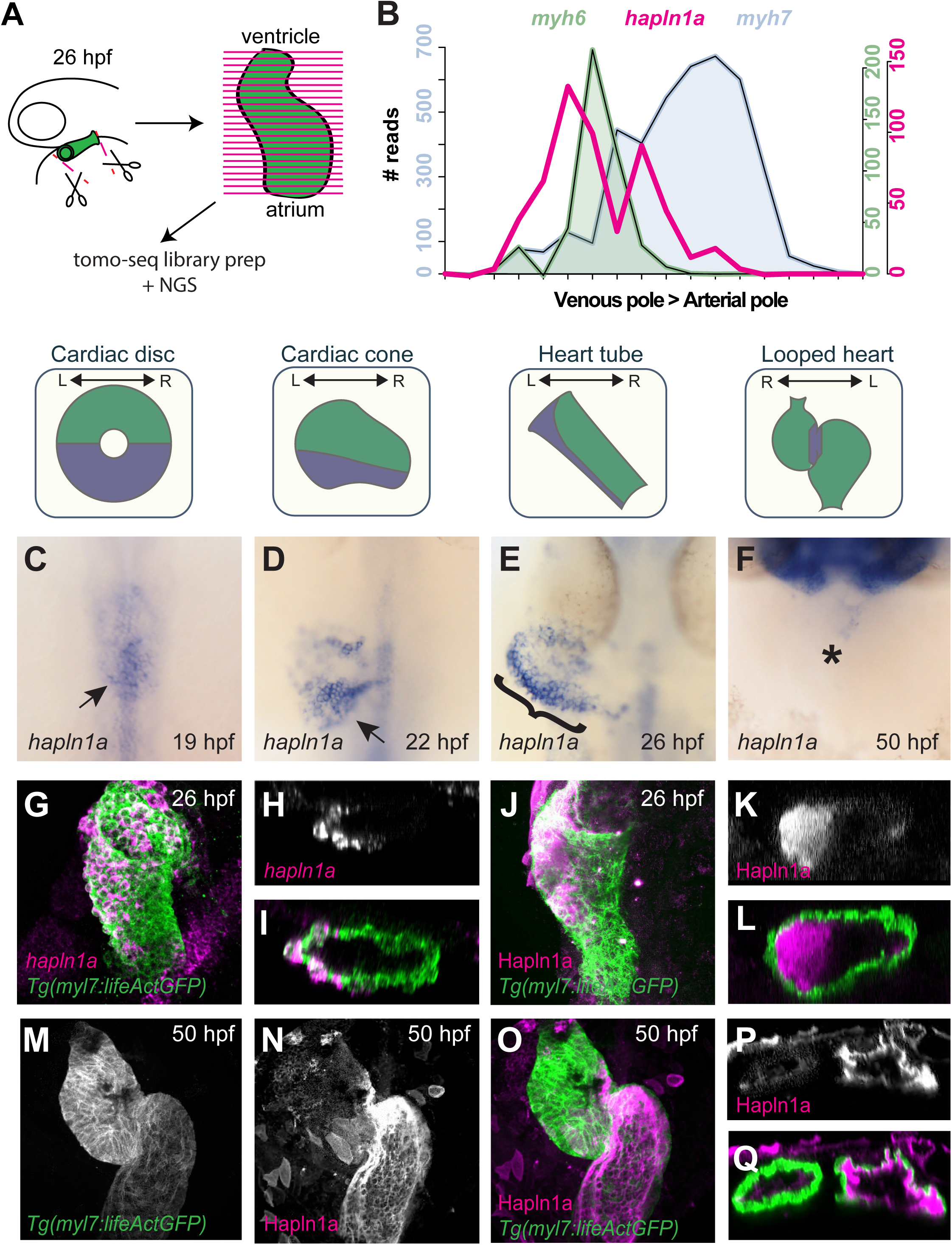
*hapln1a* is regionally expressed in the heart tube and secreted asymmetrically into the cardiac jelly. A: Schematic representation of Tomo-seq pipeline. GFP-expressing hearts are manually excised from embryos at 26hpf, and frozen in OCT tissue freezing medium. Heart tubes are sectioned along the atrioventricular axis of the heart. RNA is extracted from individual slices, labelled with a slice-specific molecular barcode during reverse transcription and undergoes RNA amplification before generating sequencing libraries. B: Example Tomo-seq traces from a single 26hpf heart tube, with individual slices from venous pole to arterial pole represented along the x axis and normalised read number plotted on the y axis. Read numbers for atrial marker *myh6* (green) and ventricular marker *myh7l* (blue) allows identification of chamber position within the dataset. *hyaluronan and proteoglycan link protein 1a* (*hapln1a*, magenta) expression is upregulated in atrial sections. C-F: mRNA *in situ* hybridisation analysis of *hapln1a* expression in the heart between 19hpf and 50hpf. At cardiac disc stage *hapln1a* is upregulated in the posterior disc (arrow C), and this posterior expression is maintained as the heart forms the cardiac cone prior to tube formation (arrow D), with lower levels found in the anterior cone. Once the heart cone has extended to form the tube, the posterior *hapln1a* expression is positioned on the left side of the tube (bracket, E), and expressed at higher levels in the atrium than the ventricle. By 50hpf *hapln1a* expression in the heart is restricted to low levels in the atrioventricular canal (AVC, asterisk, F). Schematics above *in situ* panels indicate heart morphology at each stage, and *hapln1a* expression domain within the heart. G-I: Fluorescent *in situ* hybridisation analysis of *hapln1a* (magenta) in *Tg(myl7:lifeActGFP)* transgenic embryos shows that *hapln1a* is expressed in myocardial cells at 26hpf. J-L: Fluorescent immunostaining of Hapln1a (magenta) in *Tg(myl7:lifeActGFP)* transgenic embryos demonstrates that it is secreted into the extracellular space predominantly on the left side of the heart tube (magenta) at 26hpf. G, J: dorsal views, H-I, K-L: transverse views. M-Q: Fluorescent immunostaining of Hapln1a (magenta) in *Tg(myl7:lifeActGFP)* transgenic embryos at 50hpf reveals that Hapln1a protein is maintained in the cardiac ECM as looping progresses. M-O: Ventral views, P-Q: transverse views.

Using this approach we identified *hyaluronan and proteoglycan link protein 1a (hapln1a,* formerly *crtl1)* as a candidate that might drive regionalised ECM expansion (Fig 2B). The Hapln family of proteins are secreted into the ECM where they crosslink HA to proteoglycans (Spicer et al. 2003), suggesting Hapln1a may act to modify the cardiac ECM environment. Furthermore, *Hapln1* mutant mice exhibit heart malformations including septal defects and perturbations of the inflow and outflow tracts, consistent with abnormal heart morphogenesis (Wirrig et al. 2007).

mRNA *in situ* hybridisation analysis revealed that *hapln1a* is expressed in the posterior of the heart disc and the cardiac cone (Fig 2C, D). At 26hpf *hapln1a* expression is upregulated on the left side of the cardiac tube with elevated levels of expression of *hapln1a* in the future atrium compared to the future ventricle, recapitulating the regionalised ECM expansion we observe in the heart (compare Fig 2E and Fig 1K). This is in line with recent studies demonstrating that the posterior compartment of the cardiac disc is re-positioned to the left side of the heart tube (Guerra et al. 2018). By 50hpf *hapln1a* expression is restricted to very low levels in the atrioventricular canal, the precursor to the atrioventricular valve (Fig 2F). Fluorescent *in situ* hybridization reveals *hapln1a* is expressed in the myocardium (Fig 2G-I), while analysis of Hapln1a protein localisation confirms it is deposited in the ECM (Fig 2J-L). Despite the absence of *hapln1a* expression in the heart at 50hpf (Fig 2F), Hapln1a protein is maintained in the ECM at 50hpf (Fig 2M-Q), suggesting that the ECM environment established during early stages prior to heart tube formation is maintained during heart development and may be important for continued cardiac morphogenesis.

### Hapln1a is required for heart morphogenesis and promotes ECM expansion

To determine whether Hapln1a is required for cardiac morphogenesis we used CRISPR-Cas9-mediated genome editing to generate *hapln1a* mutants. We injected a pair of guide RNAs targeting approximately 200bp upstream of the translation-initiating ATG and immediately downstream of the ATG, allowing us to excise the putative promoter of *hapln1a* (Fig S4). We recovered F1 adult fish carrying two deletions; a 187bp deletion (*hapln1a^Δ^*^187^) and a 241bp deletion (*hapln1a^Δ^*^241^), both of which remove the initiating ATG and upstream sequence, and established both as stable lines at F2. To confirm the deletions removed the *hapln1a* promoter and prevented transcription, *hapln1a* expression was analysed at 26hpf in F3 mutant embryos for each allele. Homozygous *hapln1a* promoter mutants of either allele exhibit a complete loss of *hapln1a* expression at 26hpf compared to wild type embryos (Fig 3C, E, Fig S4), demonstrating successful deletion of the *hapln1a* promoter, and confirming the promoter mutants as loss of function models. Furthermore, embryos heterozygous for a *hapln1a* promoter mutation also exhibit a reduction in levels of transcript compared to wild type siblings (Fig 3D, Fig S4). Analysis of heart development in *hapln1a^Δ^*^241^ mutants at 50hpf did not reveal striking abnormalities in cardiac morphogenesis (Fig 3F-H), however we did observe occasionally mispositioned and malformed atria. To investigate this further, we examined morphology of individual chambers at 50hpf by *in situ* hybridization analysis of the ventricular marker *myh7l* (*myosin heavy chain 7-like)* and the atrial marker *myh6* (Fig 3I-N). Quantification of either whole heart size, or individual chamber size revealed a significant reduction in whole heart size in *hapln1a^Δ^*^241^ mutants when compared to wild type siblings (Fig 3R), and a mild reduction in atrium size (Fig 3S). We observed a similar significant reduction in atrial size in the second *hapln1a^Δ^*^187^ allele (Fig S4). To quantify cardiac morphology at 50hpf we defined the looping ratio, a quotient of the looped and linear distances (Fig S4 and Methods). Although homozygous *hapln1a* mutants exhibit a reduction in cardiac size, we observed no significant reduction in looping ratio (Fig 3U, Fig S4).

**Figure 3.**
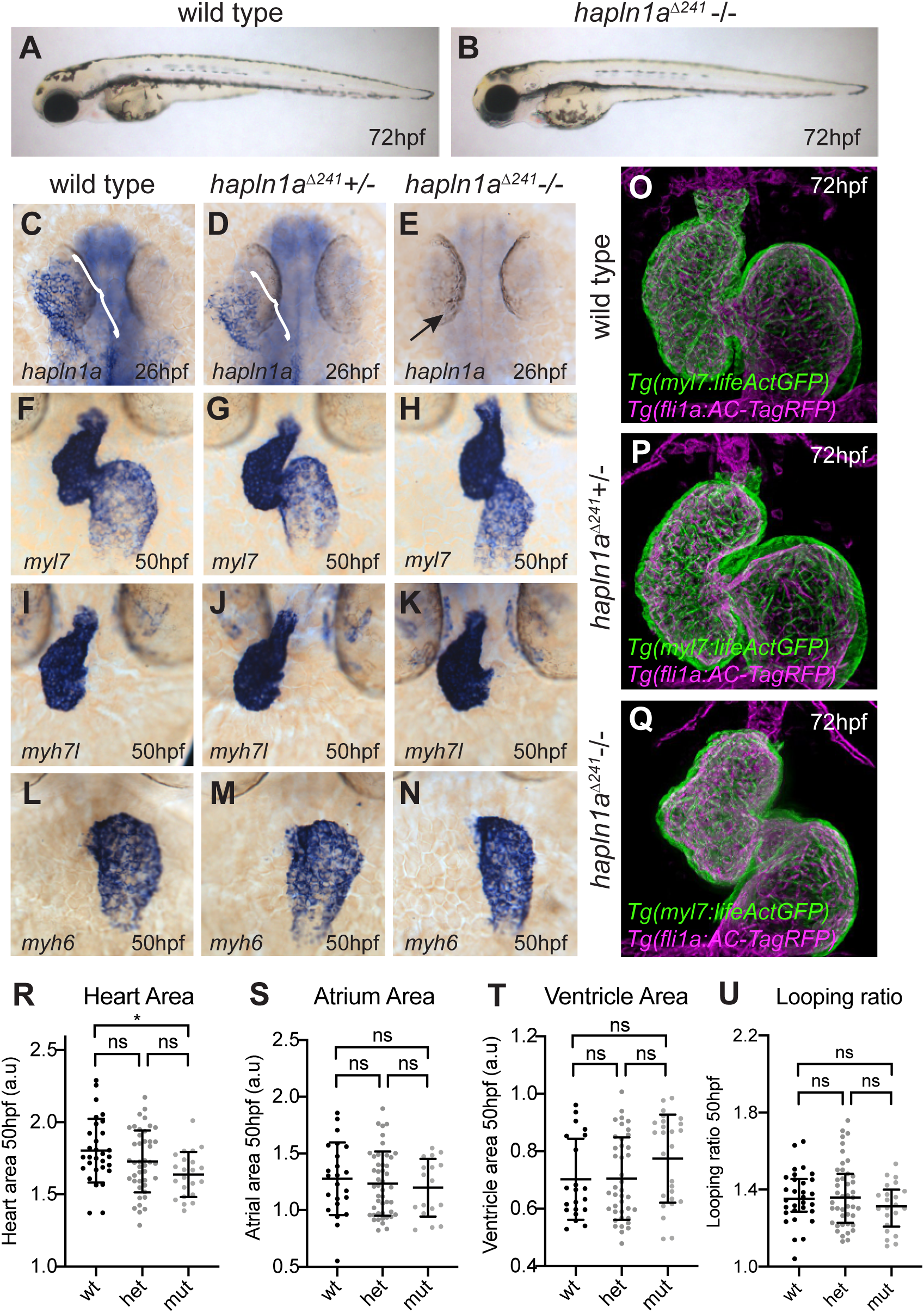
Hapln1a promotes atrial growth and heart morphogenesis. A-B : Brightfield images of wild type siblings (A) and *hapln1a^Δ241^* mutants (B) at 72hpf. C-E *hapln1a^Δ241^* mutants do not exhibit gross morphological defects. C-E: mRNA *in situ* hybridisation analysis of *hapln1a* expression at 26hpf in embryos from an incross of *hapln1a^Δ241^* heterozygous carriers. Wild type and heterozygous siblings express *hapln1a* in the heart (bracket C, D), whereas *hapln1a* is absent in homozygous mutants (arrow E). F-N: mRNA *in situ* hybridisation expression analysis at 50hpf of *myl7* (F-H), *myh7l* (I-K) and *myh6* (L-N) in wild type siblings (F, I, L), *hapln1a^Δ241^* heterozygous siblings (G, J, M) or *hapln1a^Δ241^* homozygous mutant embryos (H, K, N). O-Q: Maximum intensity projections of light sheet z-stacks of 72hpf *Tg(myl7:lifeActGFP); Tg(fli1a:AC-TagRFP)* transgenic wild type (O), *hapln1a^Δ241^* heterozygous sibling (P) and *hapln1a^Δ241^* mutant embryos (Q). R-U: Quantification of whole heart size (R) and chamber size of the ventricle (S) or atrium (T) in sibling embryos and *hapln1a^Δ241^* mutants at 50hpf. Heart size is significantly reduced and atrial size mildly reduced in *hapln1a^Δ241^* mutants compared to wild type siblings. R-T: Graphs show mean +/- SD. U: Graph shows median +/- interquartile range. * = p<0.05, ns = not significant. Comparative analysis between each group was analysed using a Kruskal-Wallis test with multiple comparisons.

To assess the impact of loss of *hapln1a* on continued morphogenesis of the heart we used live light sheet imaging of *Tg(myl7:lifeActGFP); Tg(fli1a:AC-TagRFP)* transgenic embryos, acquiring 3D images of the heart at 72hpf. We found that *hapln1a^Δ^*^241^ mutant hearts appear dysmorphic at 72hpf, with abnormally positioned atria and disrupted heart looping compared to wild type (Fig 3O-Q). Together this demonstrates that similar to mouse, *hapln1a* is required for cardiac morphogenesis.

Since Hapln1 functions as an ECM binding protein and its localisation recapitulates the regionalised ECM expansion in the heart tube prior to heart morphogenesis, we hypothesized that Hapln1a promotes cardiac morphogenesis by driving regionalised ECM expansion in the heart. Since both *hapln1a* promoter deletion alleles carry the *Tg(myl7:lifeActGFP)* transgene, this prevented analysis of ECM width throughout the heart tube of *hapln1a* mutants using the *ssNcan-GFP* HA sensor. Therefore, to examine ECM asymmetry in the heart tube upon loss of *hapln1a*, we injected a morpholino (MO) against *hapln1a* into zebrafish embryos at the 1-cell stage together with a *tp53* MO control and the *ssNcan-GFP* HA sensor and assessed ECM expansion in the heart tube at 26hpf. Analysis of Hapln1a protein levels in *hapln1a* morphants confirms successful blocking of Hapln1a translation in the morphants (Fig S5). Control embryos injected with *tp53* MO demonstrate the regionalised ECM expansion previously observed, with a left-sided expansion of the ECM (Fig 4A-B), and a higher level of ECM expansion in the atrium versus the ventricle (Fig 4C). Embryos injected with *hapln1a* MO *+ tp53* MO did not exhibit either atrial or left-sided ECM expansion (Fig 4A, B, D), suggesting that Hapln1a drives regionalised ECM expansion in the heart tube. Furthermore, light sheet imaging of *Tg(myl7:lifeActGFP); Tg(fli1a:AC-TagRFP)* transgenic embryos at 72hpf revealed that the cardiac ECM remains asymmetrically expanded in the atrium of wild type siblings (Fig 4E,G,I), but that similar to the loss of ECM asymmetry in the heart tube of *hapln1a* morphants, asymmetric ECM expansion is lost in *hapln1a* mutant hearts at 72hpf (Fig 4F,H,J). Together this supports a role for Hapln1a in regionally regulating the size of the cardiac ECM to promote normal cardiac morphogenesis.

**Figure 4.**
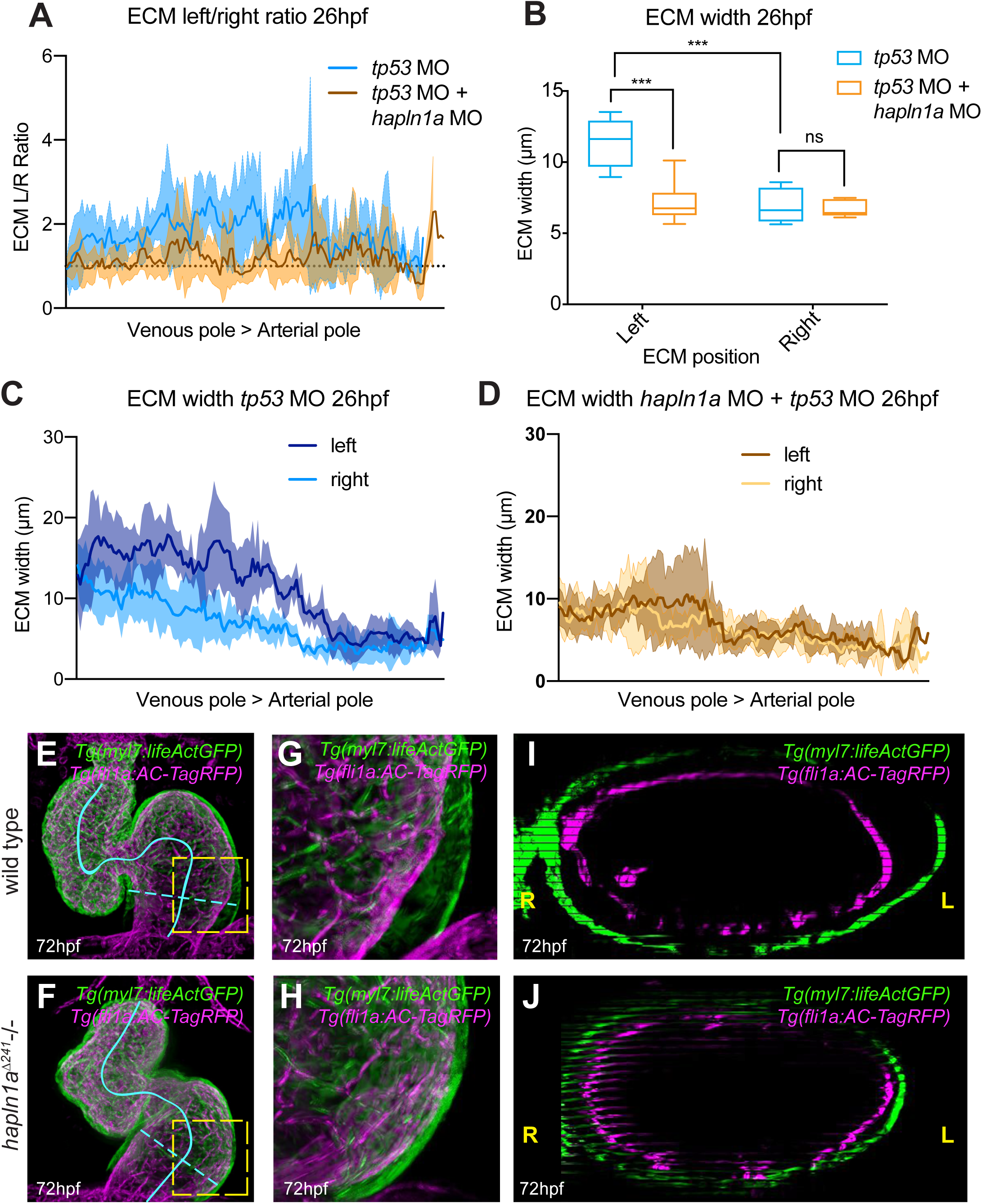
Hapln1a drives regionalised ECM expansion. A: Quantification of ECM left/right width along the longitudinal axis of the heart at 26hpf in embryos injected with either *tp53* MO (blue, n=5) or *hapln1a* MO *+ tp53* MO (orange, n=6). Mean +/- SD are plotted. B: Average ECM width on the left or right side of the heart tube in embryos injected with *tp53* MO (blue) or *hapln1a* MO *+ tp53* MO (orange). *tp53*-injected controls display a significantly expanded ECM on the left side of the heart tube compared to the right, whereas embryos injected with *hapln1a* MO *+ tp53* MO do not exhibit left-sided expansion of the ECM. Mean +/- SD are plotted. C-D: Quantification of ECM width on the left and right sides of the heart tube from venous pole to arterial pole at 26hpf in embryos injected with t*p53* MO (C) or *hapln1a* MO *+ tp53* MO (D). Graphs show mean +/- SD. The cardiac ECM in *tp53* morphants exhibits atrial and left side expansion, whereas the ECM in *hapln1a* morphants is more uniform in width from atrium to ventricle and is not expanded on the left side. Mean +/- SD are plotted, n=6. E-H: Maximum intensity projections of light sheet z-stacks of 72hpf *Tg(myl7:lifeActGFP); Tg(fli1a:AC-TagRFP)* transgenic wild type (E,G) and *hapln1a^Δ241^* mutant embryos (F,H). Yellow boxed regions (E,F) are magnified (G,H). Solid blue line indicates the centreline of the heart. I-J: Orthogonal views through the atrium of wild type (I) and *hapln1a* mutant (J) as indicated by blue dashed lines in E and F at 90° angle to the centreline. *** = p<0.001, ns = not significant. L – left, R – right.

### Hapln1a and HA interact to drive heart morphogenesis

Hapln1a is a member of a family of ECM binding proteins which crosslink hyaluronan with proteoglycans in the ECM (Spicer et al. 2003). Since *hapln1a* is transiently expressed at cardiac disc and early tube stage, this suggests that the cardiac ECM that drives continued morphogenesis of the heart is established at early stages of heart development and requires the interaction of Hapln1a with hyaluronic acid. To interrogate the temporal requirements for HA in heart looping, we applied the HA synthesis inhibitor 4-Methylumbelliferone (4-MU (Nakamura et al. 1997; Ouyang et al. 2017)) to embryos prior to the onset of heart tube formation at 18hpf, and either washed the drug off at 22hpf, or left the embryos to develop to 50hpf, when we assessed heart looping morphology. Inhibiting HA synthesis from cardiac disc stage (18hpf) until 50hpf often arrested heart development mid-way during tube formation (Fig S6), a more profound phenotype than that observed in *has2* zebrafish morphants or *Has2* mouse mutants (Smith et al. 2008; Camenisch et al. 2000). However, inhibition of HA synthesis during the short time window between cardiac disc (18hpf) and cardiac cone (22hpf) stage, prior to tube formation, resulted in normal tube formation but a specific disruption to heart looping morphogenesis (Fig S6). This supports the hypothesis that HA synthesised prior to formation of the heart tube is required for looping morphogenesis of the heart.

Having demonstrated a requirement for HA synthesis for heart morphogenesis during early cardiac development when *hapln1a* expression is initiated, we wanted to confirm the interaction of *hapln1a* and HA in heart looping morphogenesis. Injection of sub-phenotypic doses of morpholinos targeting either *has2* or *hapln1a* did not result in significant defects in cardiac morphology at 48hpf (Fig S6). However, co-injection of both *has2* and *hapln1a* morpholinos results in profound defects in heart morphology at 48hpf (Fig S6), including a reduction in heart looping ratio, and abnormal atrial morphology. This more profound phenotype than that observed by either injection of *hapln1a* MO *+ tp53* MO*, has2* MO *+ tp53* MO, 4MU treatment or deletion of the *hapln1a* promoter, suggesting that while timely HA signaling drives heart morphogenesis subsequent to tube formation, *hapln1a* is an important regional modulator of this process.

While analysis of *hapln1a* mutants demonstrates a requirement for Hapln1a in heart development (Fig 3), we wished to investigate whether the regionalization of *hapln1a* expression is important for cardiac morphogenesis. We generated a DNA construct in which the full length *hapln1a* coding sequence is driven by the pan-myocardial *myl7 (myosin light chain 7)* promoter, flanked by Tol2 transposon sites to allow integration into the genome (Fig 5A). We co-injected *myl7*:*hapln1a* DNA with *tol2* transposase mRNA at the 1-cell stage and analysed both *myl7* and *hapln1a* expression at 50hpf, allowing us to visualize heart morphology alongside assessing the extent of *hapln1a* misexpression (Fig 5B,C). We quantified the looping ratio (Fig S4), as well as the percentage coverage of *hapln1a* expression in the whole heart and plotted percentage coverage against looping ratio (Fig 5D). We found that increasing the domain of *hapln1a* expression in the heart results in a reduction in looping morphogenesis (Fig 5D), suggesting that regionalised expression of *hapln1a* in the heart is important for proper cardiac morphogenesis. Since *hapln1a* is expressed at higher levels in the atrium than the ventricle and ECM asymmetry is more robust in the atrium, we hypothesized that *hapln1a* misexpression in each chamber may impact differently on heart morphogenesis. We quantified *hapln1a* misexpression in each chamber by calculating the percentage of the chamber which expresses *hapln1a* (Fig 5E-I), and found that while misexpression of *hapln1a* in the ventricle did not impact upon heart morphogenesis (Fig 5J), misexpression of *hapln1a* in the atrium resulted in abnormal cardiac morphogenesis (Fig 5K). This suggests that spatially-restricted *hapln1a* expression in the atrium drives cardiac morphogenesis.

**Figure 5.**
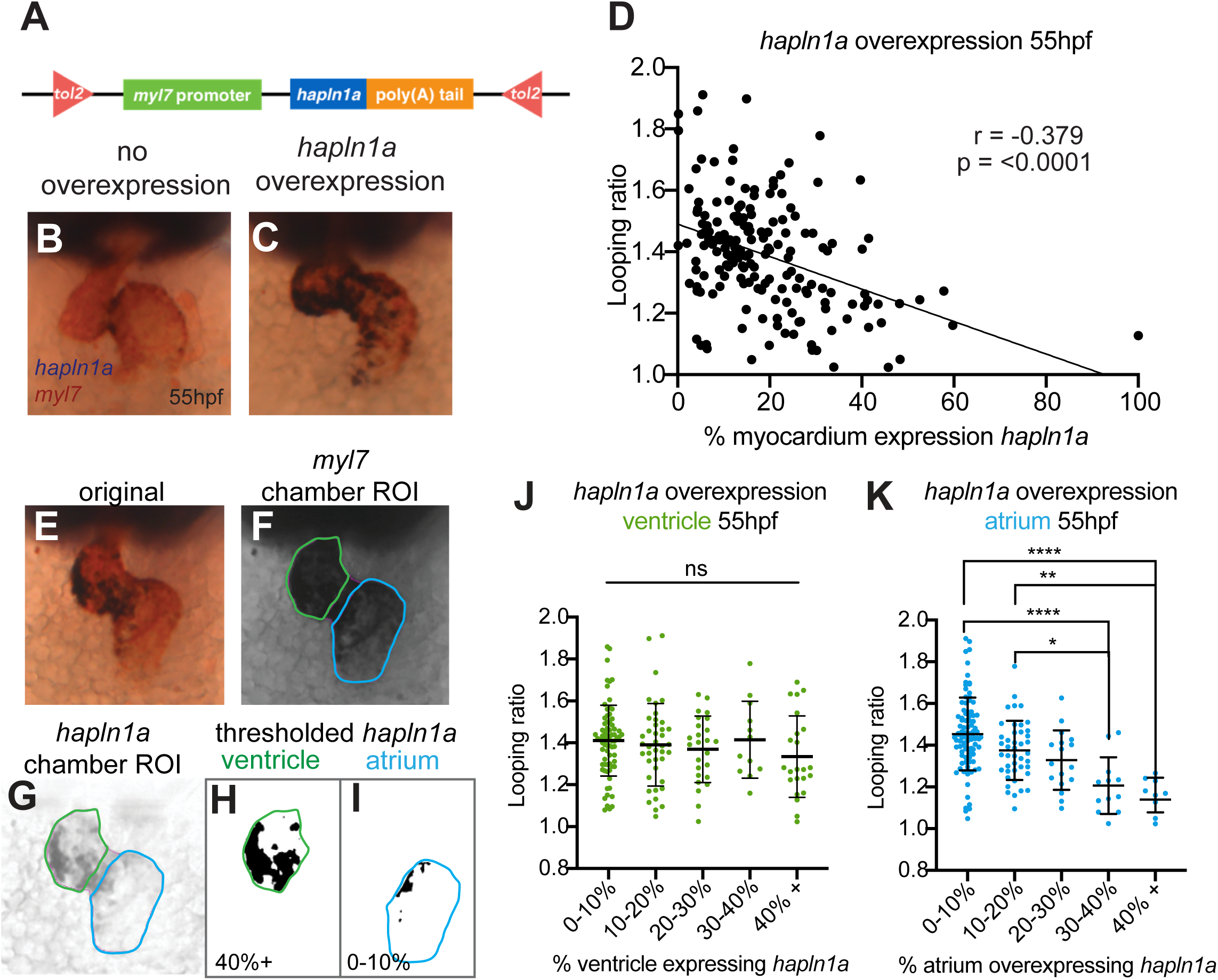
Regionalised *hapln1a* expression in the atrium promotes heart morphogenesis. A: Schematic of DNA construct used to overexpress *hapln1a* specifically in cardiomyocytes. B, C: mRNA *in situ* hybridisation analysis of *myl7* (red) and *hapln1a* (blue) at 55hpf in embryos injected with a *myl7:hapln1a* overexpression construct. D: Scatter plot depicting looping ratio as a function of percentage of the heart covered by *hapln1a* expression together with linear regression of the data (n=194). Spearman’s correlation coefficient (r) deviates significantly from zero demonstrating that increased coverage of *hapln1a* in the myocardium results in reduced heart looping morphogenesis. E-I: Illustration of quantification approach to analyse pan-cardiac or chamber-specific *hapln1a* overexpression at 55hpf. Embryos are stained for *myl7* and *hapln1a* mRNA (E), and images are split into red, blue and green channels. The green channel (F) highlights the *myl7* expression in the heart and is used to manually draw round the ventricle and atrium. The red channel highlights the *hapln1a* expression (G), and is first processed to subtract the background, before the chamber ROIs are applied (G). Each chamber ROI is isolated, the surrounding image cleared, and a threshold is applied to the *hapln1a* staining within the ROI (H-I). The number of pixels within the chamber ROI is then measured, alongside the total area of the ROI, quantifying percentage of the chamber expressing *hapln1a*. J-K: Analysis of looping ratio as a function of the level of *hapln1a* expression in each chamber of the heart. Embryos are placed into bins depending on how much of the chamber is overexpressing *hapln1a*, and average looping ratio for each bin is calculated. Overexpression of *hapln1a* in the ventricle does not affect looping ratio (J), whereas overexpression of *hapln1a* in the atrium significantly reduces looping morphogenesis (K). * = p<0.05, ** = p<0.01, **** = p<0.0001 ns = not significant.

Finally, since Hapln1a is asymmetrically expressed on the left side of the heart tube, and is required for heart morphogenesis, we hypothesized that it may contribute to a previously-described tissue-intrinsic mechanism of heart looping morphogenesis (Noël et al. 2013) and thus is expressed independent of embryonic left-right asymmetry cues. During early somitogenesis the Kupffer’s Vesicle (KV) is required to establish left-sided signalling in the embryo (Essner 2005). Embryos with mutations in *pkd2 (polycystic kidney disease 2*), which is required for KV function (Schottenfeld et al. 2007; Roxo-Rosa et al. 2015) exhibit defects in left-right asymmetry, including a disruption to normal leftward displacement of the heart tube (Schottenfeld et al. 2007). We hypothesised that induction of *hapln1a* expression occurs independent of embryonic laterality cues, but that asymmetric positioning of *hapln1a*-expressing cells in the heart tube may be tightly linked to the direction of heart tube position, and therefore dictated by embryonic left-right asymmetry. We analysed *hapln1a* expression in an incross of *pkd2^hu2173^* heterozygotes and found that consistent with our hypothesis *hapln1a* is always expressed in the posterior of the heart disc in *pkd2^hu2173^* mutants at 22hpf (Fig 6D), demonstrating that initiation of *hapln1a* expression is laterality independent. Importantly, at 26hpf we observed that positioning of *hapln1a*-expressing cells is dependent upon cardiac position – in embryos where the heart is positioned to the right, *hapln1a* is upregulated on the right side of the tube, whereas if the heart remains midline, *hapln1a* does not exhibit a clear left-right asymmetry in up-regulation (Fig 6A-C, E). These data support a model where laterality cues do not initiate *hapln1a* expression but are required for its subsequent position in the heart tube. To further investigate our model we analysed Hapln1a protein localisation in *spaw* mutant embryos, which lack asymmetric Nodal expression prior to asymmetric organ morphogenesis (Noël et al. 2013). *spaw* mutant embryos exhibit midline hearts at 26hpf and Hapln1a is no longer positioned on the left side of the heart tube as observed in sibling embryos, but instead is secreted into the cardiac ECM on the ventral face of the heart (Figure 6F-M). Together, we propose a model where initiation of *hapln1a* expression in the posterior cardiac disc is independent of KV-based laterality cues, but the subsequent cell movements which occur during heart tube formation reposition this population of cells to the left side of the heart, dictating the axis of ECM asymmetry in the heart tube (Guerra et al. 2018). Therefore, embryonic laterality positions the regionally specialized ECM in the heart tube, ensuring that directionality and growth of the heart are tightly coordinated to fine tune cardiac morphogenesis. (Figure 6N).

**Figure 6.**
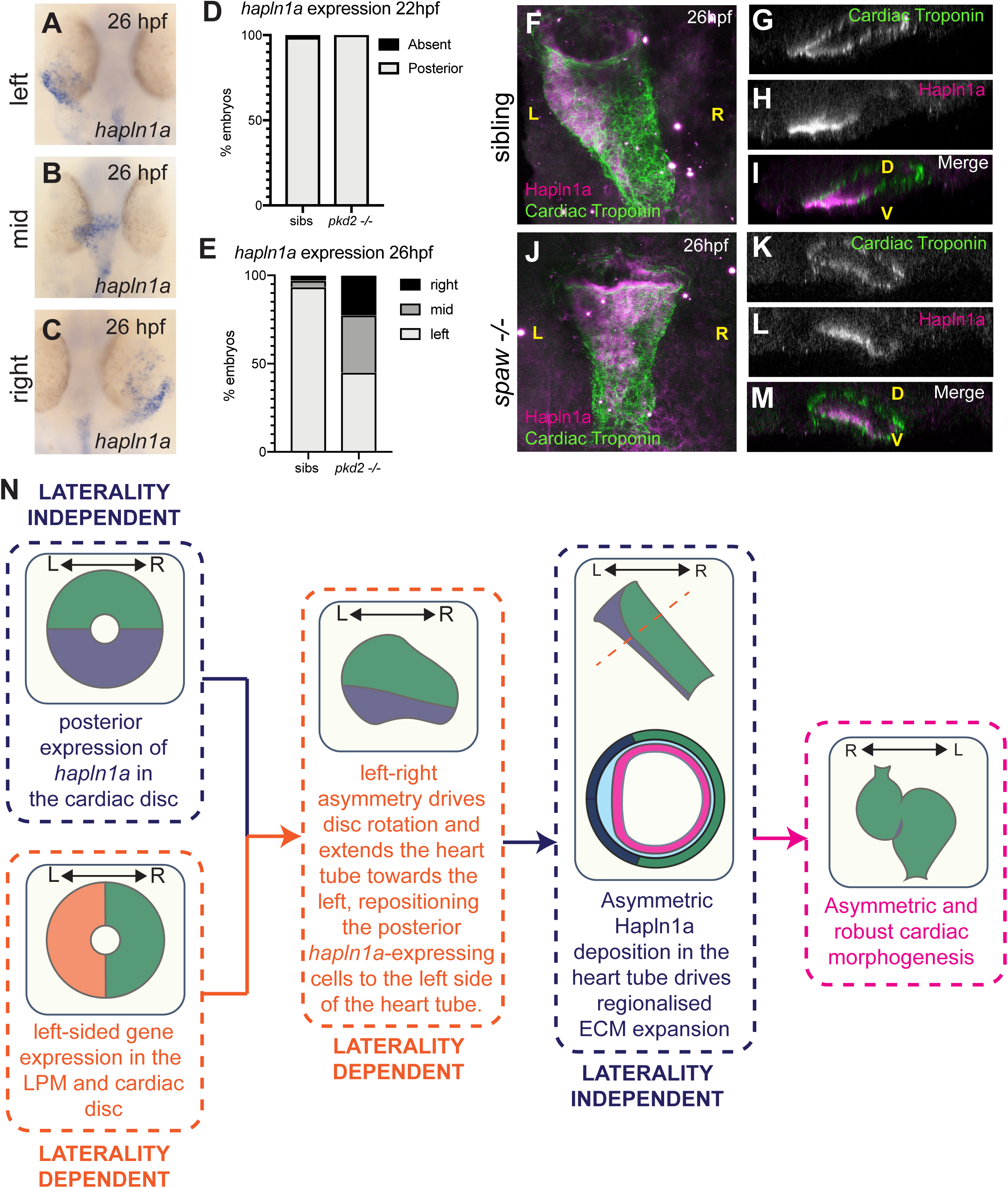
Posterior up-regulation of *hapln1a* in the cardiac disc is independent of left-right asymmetry. A-C : mRNA *in situ* hybridisation analysis of *hapln1a* expression at 26hpf in an incross of *pkd2^hu2173^* heterozygous carriers. At 26hpf, hearts that have jogged to the left exhibit left-elevation of *hapln1a* (A), hearts that remain at the midline have no clear left-right asymmetry in expression (B), and hearts on the right have right-elevated *hapln1a* (C). D-E: Quantification of position of *hapln1a* expression in sibling and *pkd2^hu2173^* mutant embryos at 22hpf (D) and 26hpf (E). F-M: Fluorescent immunostaining of Hapln1a (magenta) and cardiac troponin (green) at 26hpf in wild type siblings (F-I) or *spaw* mutant embryos (J-M). Wild type siblings exhibit left-sided deposition of Hapln1a in the heart tube (F-I, n=6), whereas *spaw* mutant embryos exhibit ventral localisation of Hapln1a (J-M, n=6). F, J: dorsal views, G-I, K-M; optical transverse sections. L - left, R - right, D - dorsal, V - ventral. N: Model depicting the interplay between laterality dependent and independent pathways that promote heart looping morphogenesis.

## Discussion

Our data show that prior to heart looping morphogenesis in zebrafish, the heart tube exhibits regionalised ECM expansion that is dependent upon localised expression of the ECM binding protein *hapln1a* and this is required to promote proper cardiac morphogenesis. Our findings build upon previous studies showing that *Hapln1* mutant mouse embryos exhibit structural cardiac malformations consistent with abnormal early cardiac morphogenesis (Wirrig et al. 2007). While that study describes expression of *Hapln1* in the valve leaflets at later stages of heart development, it does not address a potentially conserved role for transiently asymmetric *Hapln1* expression at earlier stages of heart development. Zebrafish have two *hapln1* paralogs, and each gene has a very distinct expression profile in the heart during development, with *hapln1b* being expressed primarily in the endocardium and atrioventricular canal (data not shown). Thus zebrafish provide an opportunity to define tissue-specific requirements for Hapln1 function in either the myocardium or endocardium during cardiac morphogenesis, dissecting better the specific roles for Hapln1 in cardiac development.

*hapln1a* mutants exhibit mild defects in cardiac morphology at embryonic stages, with atrial morphology predominantly affected. Interestingly, the only other gene shown to be expressed in the posterior cardiac disc/left heart tube in zebrafish is *meis2b* (Guerra et al. 2018). Analysis of *meis2b* mutants revealed defects in atrial morphology at juvenile and adult stages, supporting our conclusion that early anterior-posterior asymmetry in the heart disc/left-right asymmetry in the heart tube are important for continual cardiac morphogenesis. However, contrary to our study which reveals a reduced atrial size in *hapln1a* mutants, *meis2b* mutant adult zebrafish exhibit an enlarged atrium (Guerra et al. 2018) suggesting that while they are expressed in the same domain, these two genes regulate atrial morphogenesis differently. *hapln1a* mutants are adult viable (data not shown), therefore it would be interesting to determine whether the atrium remains underdeveloped in *hapln1a* mutants, or whether they also develop a hyperproliferative atrial hypertrophy phenotype by adulthood.

A major role of the ECM in tissue morphogenesis is to provide structural or biomechanical cues to neighbouring tissues. While *hapln1a* is expressed prior to tube formation and during very early stages of looping morphogenesis only, Hapln1a protein persists in the cardiac jelly even after the heart has undergone initial looping morphogenesis (Fig 2). Together with our data demonstrating that HA synthesis is required prior to heart tube formation to promote cardiac morphogenesis (Fig S6), this suggests that the ECM environment generated early during heart development is crucial for continual and/or maintenance of chamber morphogenesis. Interestingly, recent studies have demonstrated that Hapln1 is the key element required for tissue folding in the human neocortex, and that HA is required to maintain the architecture of the tissue after folding has occurred (Long et al. 2018). In light of this, we propose that formation of the specific ECM environment at cardiac disc stage is required to ensure the heart maintains correct shape as it undergoes early looping morphogenesis. Alternatively, Hapln1a-mediated cross-linking may modulate regional stiffness of the cardiac ECM. Differential matrix stiffness has been shown to regulate a wide variety of cellular processes which contribute to general tissue morphogenesis (Jansen et al. 2017; Wells 2008; Hannezo & Heisenberg 2019), as well as regulating cardiomyocyte form and function (Wan et al. 2019; Bhana et al. 2010; Ingber 2002; Majkut et al. 2013).

In addition to provision of mechanical cues to the surrounding cells, the ECM is also implicated in modulation of diffusion and availability of extracellular signalling molecules (Ohkawara et al. 2002; Chen et al. 2007; Müller & Schier 2011). It is therefore tempting to speculate that the specific ECM environment allows precise regionalised cellular responses to pan-cardiac or chamber specific signalling pathways.

Hapln proteins act as structural modifiers of the ECM, cross-linking HA to proteoglycans. As previously discussed, regional ECM crosslinking may change the biomechanical properties of the ECM by stabilising these components in specific regions of the heart tube. However, both HA and proteoglycan cleavage products can act as signalling molecules (Iozzo & Schaefer 2015). *Hapln1* mouse mutants exhibit a decrease in protein levels of the proteoglycan Versican (Wirrig et al. 2007), suggesting that Hapln1-mediated HA-Versican cross-linking is important for stabilising one or both of these components and preventing regionalised degradation. Further supporting an interaction between Hapln1a, Versican and HA in heart morphogenesis, both mice and medaka lacking Versican exhibit severe cardiac malformations (Mjaatvedt et al. 1998; Mittal et al. 2019), while reduction in activity of the protease ADAMTS results in reduced Versican cleavage and cardiac abnormalities (Kern et al. 2010; Kim et al. 2018). We have shown that of the two zebrafish versican paralogs, only *vcana* is expressed in a domain overlapping with *hapln1a* expression (Fig S3). Versican proteins exist in a number of isoforms and depending on domain structure can be subject to cleavage by ADAMTS proteases (Nandadasa et al. 2014). Analysis of zebrafish Vcana suggests it is a small V3 or V4-like isoform which are not predicted to undergo cleavage suggesting that in contrast to mouse, zebrafish Hapln1a may not act to stabilise Versican in the ECM. Alternatively, Hapln1a cross-linking may facilitate regional degradation of HA in the heart tube, or modulate different functions of HA, similar to the role of regionalised HA modification in establishing gut laterality in chick and mouse (Sivakumar et al. 2018). Identification of the HA receptors involved in HA signalling in the heart would help us understand how Hapln1a may regionally modulate responses to seemingly ubiquitous HA.

Recent studies have shown that cross talk between the myocardium and endocardium modulates atrial growth (Bornhorst et al. 2019), and differential ECM composition and/or degradation may help regionally fine tune this process to dictate chamber morphology. Therefore determining whether the regionalised cardiac ECM plays a structural role or is required for modulation of extracellular signalling will be an important step in understanding how these cell layers interact to drive correct chamber morphogenesis.

Analysis of *hapln1a* expression in *pkd2* mutant embryos, which have defective KV function and randomised left-right asymmetry (Schottenfeld et al. 2007) demonstrates that posterior up-regulation of *hapln1a* in the cardiac disc is not dependent upon KV function. However, we observe that in *pkd2* mutants where asymmetry of the heart is reversed and the heart tube extends to the right, due to the cellular movements required for tube formation *hapln1a*-expressing cells are positioned on the right side of the heart tube instead of the left (Fig. 6). We propose a model in which while antero-posterior patterning of the heart disc is laterality-independent, since laterality signals promote cardiac disc rotation and heart tube displacement, this results in *hapln1a* asymmetry along the left-right axis of the heart tube. This generates lateral ECM asymmetry in the heart, promoting robust looping and chamber ballooning morphogenesis (Figure 6N). This would begin to explain previous observations that heart tube position prior to looping predicts the direction of looping morphogenesis (Baker et al. 2008; Chen et al. 1997), since the direction of heart tube extension will dictate lateralised ECM asymmetry in the tube. In addition, we show that in embryos lacking Nodal signalling, Hapln1a is positioned on the ventral face of the heart (Fig 6). Together with our previous observations that the heart disc of *spaw* mutants undergo a very mild level of rotation and that the direction of this limited rotation is consistent with the final outcome of looping direction (Noël et al. 2013), it is possible that even slight rotation in the heart disc is sufficient to set up mild asymmetry in the ECM that can go on to promote directional looping, supporting the hypothesis that these pathways may act together to robustly ensure directional looping morphogenesis.

Together this study elaborates upon our previously proposed model where laterality-based extrinsic cues feed into a tissue-intrinsic mechanism of heart looping to promote robust directional cardiac morphogenesis.

## Acknowledgements

The Zeiss Z1 Lightsheet microscope was funded by BHF Infrastructure grant IG/15/1/31328. Additional imaging work was performed at the Wolfson Light Microscopy Facility, using the Airyscan and Nikon A1 microscopes for acquisition and Arivis Vision4D for image processing. EN is supported by British Heart Foundation Intermediate Basic Science Research Fellowship grant number FS/16/37/32347, and an Academy of Medical Science Springboard Award. We thank Kelly Smith for the *ssNcan-GFP* construct, Markus Affolter for the VE-Cadherin antibody, and Aylin Metzner for help with characterising the *pkd2^hu2173^* allele. We also thank Tanya Whitfield, David Strutt and Simon Johnston for their helpful comments on the manuscript.

## Author Contributions

CJD and EN conceived the study and designed the experiments. CJD, JSP, FH, EP and EN carried out experimental work. AS, RW and TC shared the *Tg(fli1a:AcTagRFP)* transgenic line prior to publication, FT and JB generated the *Tg(lft2BAC:Gal4FF)* transgenic line, and FvE characterised the *pkd^hu2173^* allele. JB provided financial support for the Tomo-seq experiments. EN wrote the manuscript with input from CJD, JSP, EP, FT and TC.

## Materials and Methods

### Zebrafish maintenance

Adult zebrafish were maintained according to standard laboratory conditions. The following lines were used: AB, Tg(myl7:eGFP) (Huang et al. 2003), Tg(myl7:lifeActGFP) (Reischauer et al. 2014), Tg(fli1a:AC-TagRFP)^sh511^ (Savage et al. 2019), spaw^t30973^ (Noël et al. 2013), Tg(lft2BAC:Gal4FF); Tg(UAS;RFP), pkd2^hu2173^, hapln1a^promΔ241^ (allele designation hapln1a^sh580^), hapln1a^promΔ187^(allele designation hapln1a^sh578^). Embryos older than 24hpf were treated with 0.2 mM 1-phenyl-2-thiourea (PTU) in E3 medium to inhibit melanin production.

### Generation of hapln1a mutants

To generate hapln1a mutant zebrafish lines, CRISPR guide RNAs (gRNA) were designed to target the putative promoter region of hapln1a (GRCz11: ENSDART00000122966.4, g1: 5’-TCGTCTCTCTCTAAGGGGAGGGG-3’) and the downstream region of the translation start site (g2: 5’-GATGATTGCTCTGTTTTCTGTGG-3’). Sequence-specific CRISPR RNAs (crRNA) were synthesised by Merck, resuspended in MilliQ water to 21.4μM, and injected together with an equimolar concentration of trans-activating RNA (tracrRNA, Merck) and Cas9 protein (NEB M0386T) into the yolk of 1-cell stage embryos in a volume of 1nl. CRISPR-Cas9-injected embryos were raised to adulthood (F0) and individuals transmitting putative promoter deletions in the germline were identified by outcrossing to wildtype. Embryos collected from these outcrosses were genotyped by PCR using the following primers to amplify the putative promoter region of hapln1a: forward 5’-ACATTTTGCATGCCCTCGAA-3’; reverse 5’-TGCATCCTGGACCTTCATTCA-3’. Successful promoter deletion was identified by presence of a smaller PCR fragment and subsequent Sanger sequencing of the promoter region to confirm the deletion. F0 founders transmitting a desirable mutation were outcrossed, offspring raised to adulthood, and heterozygous F1 adults identified by genotyping using the above primers. Two separate hapln1a promoter deletion alleles were recovered: hapln1a^promΔ187^ and hapln1a^promΔ241^.

### Generation of the Tg(lft2BAC:Gal4FF) transgenic line

The Tg(lft2BAC:Gal4FF) line was generated by recombineering of bacterial artificial chromosome (BAC) CH211-236P5 as previously described(Bussmann & Schulte-Merker 2011; Tessadori et al. 2012). A Gal4FF_kan cassette was inserted at the ATG start codon of the first exon of the lft2 gene. Amplification from a pCS2+Gal4FF_kanR plasmid was achieved with primers : F_LFT2_GAL4FF 5’-cctcagagcttcagtcagtcattcattctttcactggcatcgttagatcaACCATGAAGCTACTGTCTTCTATCGA AC-3’ R_LFT2_NEO 5’-tgtgtgagtgagatcgctgtggtcaaaatgaacagctggatgaacagagcTCAGAAGAACTCGTCAAGAAGGC G-3’ Sequences homologous to the genomic locus in lower case. Recombineering was essentially carried out following the manufacturer’s protocol (Red/ET recombination; Gene Bridges GmbH). BAC DNA isolation was carried out using a Midiprep kit (Life Technologies BV). BAC DNA was injected at a concentration of 300 ng/µl in the presence of 0.75U PI-SceI meganuclease (New England Biolabs) in 1-cell stage Tg(UAS:GFP) or Tg(UAS:RFP) embryos (both UAS lines(Asakawa & Kawakami 2008)). At 1dpf, healthy embryos displaying robust lft2-specific fluorescence were selected and grown to adulthood. Founder fish (F0) were identified by outcrossing and the progeny (F1) was grown to establish the transgenic line.

### Generation of pkd2^hu2173^ allele

The pkd2^hu2173^ allele was generated using ENU mutagenesis and consists of an A->T transversion at base position 1327 which results in a premature stop codon at amino acid 302 of 904. The truncation occurs in the first extracellular loop, before the channel pore, and is predicted to be a null. The pkd2^hu273^ allele can be identified by PCR amplification with the following primers: forward primer 5’-GATTTATTGCTCTGTTTGTTTGTAAGGA-3’ and reverse primer 5’ -GAAGTCCAAGAACACCGCTC-3’, followed by XmnI restriction of the PCR product. The primers contain a mismatch which together with the pkd2^hu2173^ mutation introduces an XmnI recognition site into the mutant strand.

### mRNA in situ hybridisation

Embryos were fixed overnight in 4% PFA, and mRNA in situ hybridisations were carried out as previously described (Noël et al. 2013). Fluorescent in situ hybridisations were performed using the TSA kit (Perkin-Elmer) (Welten et al. 2006). The hapln1a mRNA probe construct was generated by amplifying an 860bp fragment of the hapln1a CDS using the primers F: 5’-TGGCATTGATGGTGTTTGCA-3’; R: 5’-ACAGTTCCGTCACTAAGCCA-3’. The has2 mRNA probe construct was generated by amplifying an 1050bp fragment of the CDS using the primers F: 5’-GTTCACGCAGACCTCATCAC-3’; R: 5’-CATCCAATACCTCACGCTGC-3’. The acana mRNA probe construct was generated by amplifying an 1067bp fragment of the CDS using the primers F: 5’-CGGATCAAGTGGAGTCTGGT-3’; R: 5’-GAAGGGAGGACGTGGGAAAT-3’. The acanb mRNA probe construct was generated by amplifying an 1035bp fragment of the CDS using the primers F: 5’-ATCAAGACAGCACCCTCAGT-3’; R: 5’-TTTCTGGAAATGGCGTGGTC-3’. The chsy1 mRNA probe construct was generated by amplifying an 801bp fragment of the CDS using the primers F: 5’-CACCATTCAGCTCCATCGTG-3’; R: 5’-TCGGCTTTGGGGTACTTCAT-3’. All probe sequences were ligated into the PCR2-TOPO vector (Invitrogen). vcana and vcanb mRNA probes have been previously described (Kang et al. 2004). Riboprobes were transcribed from linearized template in the presence of DIG-11-UTP or Fluorescein-11-UTP (Roche).

### Immunohistochemistry

Whole mount immunohistochemistry was carried out as previously described (de Pater et al. 2009). The following commercially available primary antibodies were used: αGFP (1:1000 Aves lab), αCT3 (1:100, Developmental Studies Hybridoma Bank), αCdh5 (Blum et al. 2008) (1:100). A rabbit polyclonal antibody targeting amino acids 117-134 (DGMNDMTLEVDLEVQGKD) of zebrafish Hapln1a was designed and produced by Proteintech. Test bleeds were used to determine cross-reactivity with Hapln1a by comparing protein localisation at 26hpf with mRNA in situ hybridization. Subsequently, affinity-purified Hapln1a antibody was used 1:100. Fluorophore-conjugated secondary antibodies were obtained from Jackson labs and used at 1:200.

### Tomo-seq

Hearts were dissected from Tg(myl7:eGFP) zebrafish embryos at 26hpf and placed into O.C.T cryofreezing medium (Sakura Finetek). Blue Affy-gel beads (BioRad) were placed at each end of the heart tube to aid visualisation during sectioning, and the hearts were rapidly frozen in OCT blocks and stored at −80°C. Hearts were sectioned using a cryostat at 9nM resolution. RNA extraction, aRNA synthesis, library preparation, sequencing and data analysis was performed as previously described (Burkhard & Bakkers 2018; Junker et al. 2014).

### Morpholino-mediated knockdown and hapln1a overexpression analysis

The following morpholino was designed to target the translational start site of hapln1a (AGAGCAAT[CAT]CTTCACGTTTGTTA). Morpholinos blocking tp53 (Robu et al. 2007; Langheinrich et al. 2002), Zfin tp53 MO-4) and has2 (Bakkers et al. 2004), has2 MO-1) are previously described. All morpholinos were supplied by GeneTools and diluted to a stock of 1mM. Working concentrations were as follows: hapln1a 500nM or 250nM, has2 250nM, combinatorial has2/hapln1a 250nM each, tp53 250nM. All has2 and hapln1a morpholinos were co-injected together with the tp53 morpholino. Embryos were injected with 1nl of working morpholino solution.

Full length hapln1a coding sequence was amplified using the following primers containing AttB sequences for subsequent Gateway cloning, as well as a Kozak sequence (underlined): F: 5’ggggacaagtttgtacaaaaaagcaggct<u>TCGCCGCCACC</u>ATGATTGCTCTGTTTTCTGT 3’; R: 5’GGGGACCACTTTGTACAAGAAAGCTGGGTTTTACTGCTGGGCTTTGTAGCAATA 3’. The resulting PCR product was ligated into the pDONR221 middle entry Gateway vector, and sequenced to verify integrity of the insertion, generating a pMEhapln1aCDS vector. Full length hapln1a was subsequently recombined with a p5E myl7 promoter sequence, and a p3E polyA sequence into the pDestTol2pA3 destination vector to generate the final pDestmyl7:hapln1a overexpression construct. All Gateway cloning was carried using the Tol2 kit via standard protocols (Kwan et al. 2007). 60pg of pDestmyl7:hapln1a was co-injected with 25pg of tol2 mRNA into the cell of 1-cell stage embryos. Analysis of hapln1a overexpression and cardiac morphogenesis was carried out using double in situ hybridisation to assess hapln1a and myl7 expression.

### RNA injections

ssNcan-GFP mRNA was synthesised from the ssNcan-GFP construct as previously described (De Angelis et al. 2017). Embryos were injected with 100pg of mRNA in 1nl volume at the 1-cell stage and screened for GFP at 24hpf.

### Pharmacological treatments

To block HA synthesis, 4-Methylumbelliferone (4-MU, Sigma-Aldrich) was dissolved in DMSO to a stock concentration of 100mM, and subsequently diluted to a working concentration of 1mM in E3 medium. Embryos were dechorionated and incubated in 4-MU or an equal concentration of DMSO (1%). For timed treatments, at the end of the treatment window embryos were washed 3 x 5 mins in E3 to remove the 4MU, before being placed in fresh E3 until fixation.

### Imaging and image quantification

Live zebrafish embryos were imaged on a ZEISS Lightsheet Z.1 microscope at 72hpf. To assess cardiac morphology at 72hpf embryos were anesthetised with tricaine before mounting in 1% low melting point agarose in E3 with 8.4% tricaine, using black capillaries. To stop the heart the imaging chamber was filled with E3 plus tricaine (8.4%) and the temperature maintained at 10°C. All samples were imaged using a 20X lens and 1.0 zoom. Dual side lasers with dual side fusion and pivot scan were used for sample illumination. Image stacks were initially processed using Vision4D (Arivis AG, Germany) and Fiji. Processing steps included noise removal, background correction, and subsequent application of individual morphological filters to each channel to sharpen the edges of the myocardial and endocardial tissue layers. Maximum intensity z-projections of the composite channels were used to visualise cardiac morphology the centreline of the heart was manually traced. Optical transverse sections perpendicular to the centreline were generated at regularly-spaced intervals originating from the venous pole into the atrium to visualise the cardiac ECM.

Embryos injected with ssNcan-GFP mRNA were fixed overnight in 4% PFA, and immunohistochemistry was carried out to amplify the GFP signal. Embryos were dissected and imaged using a Zeiss Airyscan microscope, and z stacks were obtained with a z-step size of 1µM. Images were Airyscan processed using Zen Black software (Zeiss), and the resulting image stacks were optically resliced using Fiji. ECM width was manually measured in Fiji. ECM measurements were aligned between samples at the venous pole of the heart for plotting. Looping ratio was calculated from images of myl7 expression detected by ISH. All samples from one experimental set were blinded using the ImageJ Blind_Analysis plugin (https://github.com/quantixed/imagej-macros/blob/master/Blind_Analysis.ijm). The linear distance from arterial to venous poles of the heart was measured as a straight-line distance, and looped distance was drawn from the same positions at each pole through the centre of each chamber, down the centreline of the looped heart. Looping ratio was determined by dividing looped distance by linear distance. Statistical testing of average looping ratio between experimental conditions was carried out using Kruskal-Wallis with Dunn’s multiple comparisons.

Heart, ventricle or atrium area at 50hpf was quantified from in situ hybridisations by manually drawing round either myl7, myh7l or myh6 staining area in Fiji. Statistical testing of heart or chamber size between genotypes was carried out using Kruskal-Wallis with Dunn’s multiple comparisons.

Quantification of hapln1a overexpression was performed by imaging overexpression embryos where myl7 expression was detected using INT/BCIP and hapln1a expression detected using NBT/BCIP. All embryos were imaged using the same microscope settings, and individual images combined into a composite of all experiments. Using the composite, channels were split in Fiji, resulting in the blue and green channels carrying the myl7 stain, and the red channel carrying the hapln1a stain. Background was subtracted in the red channel. The myl7 staining was used to calculate looping ratio for each heart. In addition, each myl7 signal was used to manually trace the whole heart, atrium, AVC and ventricle for each heart, which was saved as a region of interest (ROI) and the area of each chamber was measured in pixels. Next, the hapln1a image was thresholded to generate a binary image. The whole-heart, or chamber-specific ROI was applied to the thresholded hapln1a channel, and the number of positive pixels in each ROI recorded. Number of positive pixels are as a % of total number of pixels comprising the heart or the specific chamber could then be calculated and plotted against looping ratio for each heart. Spearmann’s correlation coefficient (r) was calculated in GraphPad Prism, with 95% confidence intervals.

