## Supplemental Figures and Legends for "Asymmetric Hapln1a drives regionalised cardiac ECM expansion and promotes heart morphogenesis during zebrafish development"

**Derrick *et al* 2019 Supplemental Data**

**Figure S1 – Generation of the *Tg(lft2BAC:Gal4FF)* transgenic zebrafish**

Characterisation of the *Tg(lft2BAC:Gal4FF)* reporter line. (A) An expression cassette containing Gal4FF was recombined into BAC CH211-236P5 at the ATG site of the 1st exon of the *lft2* gene. (B-G) GFP expression pattern of the *Tg(lft2BAC:Gal4FF)*; *Tg(UAS:GFP)* double transgenic embryos at 80% epiboly (B-E) and *in situ* hybridisation analysis of *gfp* mRNA (F,G). (H-I) mRNA *in situ* hybridisation analysis of *lft2* expression at 80% epiboly. Note the overlap of expression at the forming midline (compare D, G, I). (J-P) Reporter expression (RFP) in *Tg(lft2BAC:Gal4FF); Tg(UAS:RFP*); *Tg(myl7:eGFP)* triple transgenic embryos at 22 somites (lateral view, J) and in the cardiac region at 28 hpf (lateral view, K-M) and at 50 hpf (frontal view, N-P).

**Figure S2 – Cardiac jelly in the atrium is asymmetric at 50hpf**

A: Schematic representing the positioning of the heart in a zebrafish embryo at 50hpf (ventral view). Dotted line demarcates the optical transverse sections imaged in panels B-G. B-G: Light sheet optical cross-sections through the atrium of a 50hpf *Tg(myl7:lifeActGFP); Tg(fli1a:AC-TagRFP); Tg(lft2BAC:Gal4FF); Tg(UAS:RFP*) transgenic embryo during diastole (B-D) and systole (E-G). The myocardium is highlighted in green (B, D, E, G), and the dorsal myocardium and endocardium are highlighted in magenta (C, D, F, G). The extracellular space between the myocardium and endocardium is expanded on the left side of the atrium (white arrowhead). H: Quantification of left-right ECM ratio in the atrium at 50hpf, where a value greater than 1 (red dotted line) denotes a left-sided expansion (n=6).

**Figure S3 – The ECM constituents HA, *versican* and *aggrecan,* and the ECM synthases *has2* and *chsy1* are not asymmetrically expressed in the heart tube**

A: Schematic representing the positioning of the heart in a zebrafish embryos at 28hpf (dorsal view), with the dotted line representing the optical transverse sections imaged in panels B-G. B-G: Light sheet optical cross-sections through the heart tube of a 26hpf *Tg(fli1a:AC-TagRFP); Tg(lft2BAC:GalFF); Tg(UAS:RFP)* transgenic embryo injected with *ssNcan-GFP* RNA during diastole (B-D) and systole (E-G). The *ssNcan-GFP* sensor is highlighted in green (C, D, F, G), and the dorsal myocardium and endocardium are highlighted in magenta (B, D, E, G). The *ssNcan-GFP* binds to HA and is found in the extracellular space between the myocardium and endocardium and does not appear to display asymmetric distribution. L – left, R – right. H-S: mRNA *in situ* hybridisation analysis of *has2* (H, I), *vcana* (J, K), *vcanb* (L, M), *chsy1* (N, O), *acana* (P, Q), and *acanb* (R, S) at 21hpf and 26hpf. None of the genes analysed display anterior-posterior asymmetry in the heart disc or left-right asymmetry in the heart tube. Dorsal views, anterior to top.

**Figure S4 – Hapln1a is required for cardiac morphogenesis**

A: Schematic of deletion of putative promoter in *hapln1a* mutants by CRISPR-Cas9 mediated genome editing, based on Ensembl GRCz11 *hapln1a* transcript ID ENSDART00000122966.4. B-D: mRNA *in situ* hybridisation analysis of *hapln1a* expression at 26hpf in embryos from an incross of *hapln1a*$^{\Delta 187}$ heterozygous carriers. Wild type and heterozygous siblings express *hapln1a* in the heart (brackets, B, C), whereas *hapln1a* is absent in homozygous mutants (arrow D). E-M: mRNA *in situ* hybridisation expression analysis at 50hpf of *myl7* (E-G), *myh7l* (H-J) and *myh6* (K-M) in wild type siblings, *hapln1a*$^{\Delta 187}$ heterozygous siblings or *hapln1a*$^{\Delta 187}$ homozygous mutant embryos. (N-P) Illustration of technique used to quantify heart looping ratio. mRNA *in situ* hybridisation analysis of *myl7* expression is used to demarcate the heart.

The straight-line distance is measured (white line, N). A second line is drawn from the same position at the poles following the midline of the chambers of the heart (the centreline, green line O). Looping ratio is a quotient of the looped and linear distances (P). Q-T: Quantification of heart size (Q), or chamber size of the ventricle (R) and atrium (S) in sibling embryos and *hapln1a*$^{\Delta 187}$ mutants at 50hpf. Atrial size is reduced in *hapln1a*$^{\Delta 187}$ mutants compared to wild type siblings (S). Mean +/- SD are plotted. T: Quantification of looping ratio in sibling embryos and *hapln1a*$^{\Delta 187}$ mutants at 50hpf. Median +/- interquartile range is plotted * = $p<0.05$, ns = not significant.

**Figure S5 – *hapln1a* morpholino is a loss-of-function model and recapitulates *hapln1a* mutant cardiac phenotypes**

A-F: Immunohistochemistry analysis of Hapln1a (magenta) in embryos injected with either *tp53* MO (A-C) or *hapln1a MO* + *tp53 MO* (D-F). Cardiac Troponin (green) outlines the heart. Hapln1a is expressed in *tp53* MO-injected embryos (n=5) but is absent in embryos injected with *hapln1a MO* + *tp53 MO* (n=6). G-I: mRNA *in situ* hybridisation analysis of *myl7* expression at 50hpf in uninjected controls (G), embryos injected with *tp53* MO (H) or *hapln1a* MO + *tp53* MO (I).

**Figure S6 – HA is required at early stages of heart development for cardiac morphogenesis and interacts with Hapln1a**

A: Schematic depicting timing of 4-MU treatments. B-F: *in situ* hybridisation analysis of *myl7* expression at 48hpf in embryos treated either with DMSO (B, E) or 4-MU (C, F). Embryos treated with 4-MU from 18hpf to 48hpf exhibit severe defects in heart development, characterised by a failure to correctly form the heart tube (C). In comparison, embryos treated with 4-MU in a shorter time window from 18hpf to 22hpf exhibit milder defects in heart morphogenesis, characterised a specific failure in heart looping morphogenesis (F). G-K: mRNA *in situ* hybridisation analysis at 48hpf of *myl7* expression in embryos injected with *tp53* MO (H), sub-phenotypic doses of either *has2* MO + *tp53* MO (I) or *hapln1a* MO + *tp53* MO (J), or a co-injection of sub-phenotypic doses of both *has2* and *hapln1a* MOs together with *tp53* MO (K). Embryos injected with sub-phenotypic doses of either *has2* MO + *tp53* MO or *hapln1a* MO + *tp53* MO do not exhibit heart defects, whereas embryos co-injected with both morpholinos together with *tp53* MO display unlooped hearts with dilated atria (K). L: Quantification of heart looping ratio at 48hpf in embryos injected with *tp53* MO, subphenotypic doses of single *has2* or *hapln1a* MO, or coinjection of *has2* and *hapln1a* MOs. Embryos coinjected with *has2* and *hapln1a* MOs exhibit significantly reduced heart looping compared. Median +/- interquartile range are plotted * = $p<0.05$, ** = $p<0.01$, *** = $p<0.001$, **** = $p<0.0001$.

**Movie S1**

Live light sheet optical cross-sections through the beating heart tube of a 26hpf *Tg(myl7:lifeActGFP); Tg(fli1a:AC-TagRFP) ;Tg(lft2BAC:Gal4FF,UAS:RFP)* at the level of the future atrium. The myocardium is marked in green and the dorsal myocardium and endocardium are marked in magenta. Movie speed is not in real-time.

**Table S1: Number of embryos analysed for *hapln1a* morphant, *hapln1a* mutant and *pkd2* mutant experiments**

| Experiment | Stage/gene | Uninjected | *tp53* MO | *tp53* MO + *hapln1a* MO |
|---|---|---|---|---|
| *hapln1a* MO | 50hpf/*myl7* | 72 | 21 | 111 |

| Genotype | Stage/gene | Wild type | Heterozygous | Homozygous mutants |
|---|---|---|---|---|
| *hapln1a*$^{\Delta 187}$ mutants | 50hpf/*myl7* | 28 | 40 | 15 |

| | | | | |
|---|---|---|---|---|
| | 50hpf/*myh7l* | 29 | 51 | 17 |
| | 50hpf/*myh6* | 25 | 50 | 30 |
| *hapln1a*$^{\Delta 241}$ mutants | 50hpf/*myl7* | 30 | 43 | 22 |
| | 50hpf/*myh7l* | 21 | 40 | 26 |
| | 50hpf/*myh6* | 23 | 46 | 17 |
| *pkd2* $^{hu2173}$ mutants | 22hpf/*hapln1a* | 72 | | 13 |
| | 26hpf/*hapln1a* | 138 | | 40 |

**Table S2: Raw Tomo-seq transcriptional analysis of Heart #1 at 26hpf**

Heart 1 sectioned from venous pole to arterial pole, table includes Ensembl gene identifiers, raw read numbers per section, and ERCC spike in read numbers

**Table S3: Spike-in normalised transcriptional analysis of Heart #1 at 26hpf**

Heart 1 sectioned from venous pole to arterial pole, read numbers per section normalised to spike-in RNA, and including gene names.

**Table S4: Raw Tomo-seq transcriptional analysis of Heart #2 at 26hpf**

Heart 1 sectioned from arterial pole to venous pole, table includes Ensembl gene identifiers, raw read numbers per section, and ERCC spike in read numbers

**Table S4: Spike-in normalised transcriptional analysis of Heart #2 at 26hpf**

Heart 2 sectioned from venous pole to arterial pole, read numbers per section normalised to spike-in RNA, and including gene names.

Supplementary Figure 1 - Generation of *Tg(lft2BAC:Gal4FF)* transgenic zebrafish

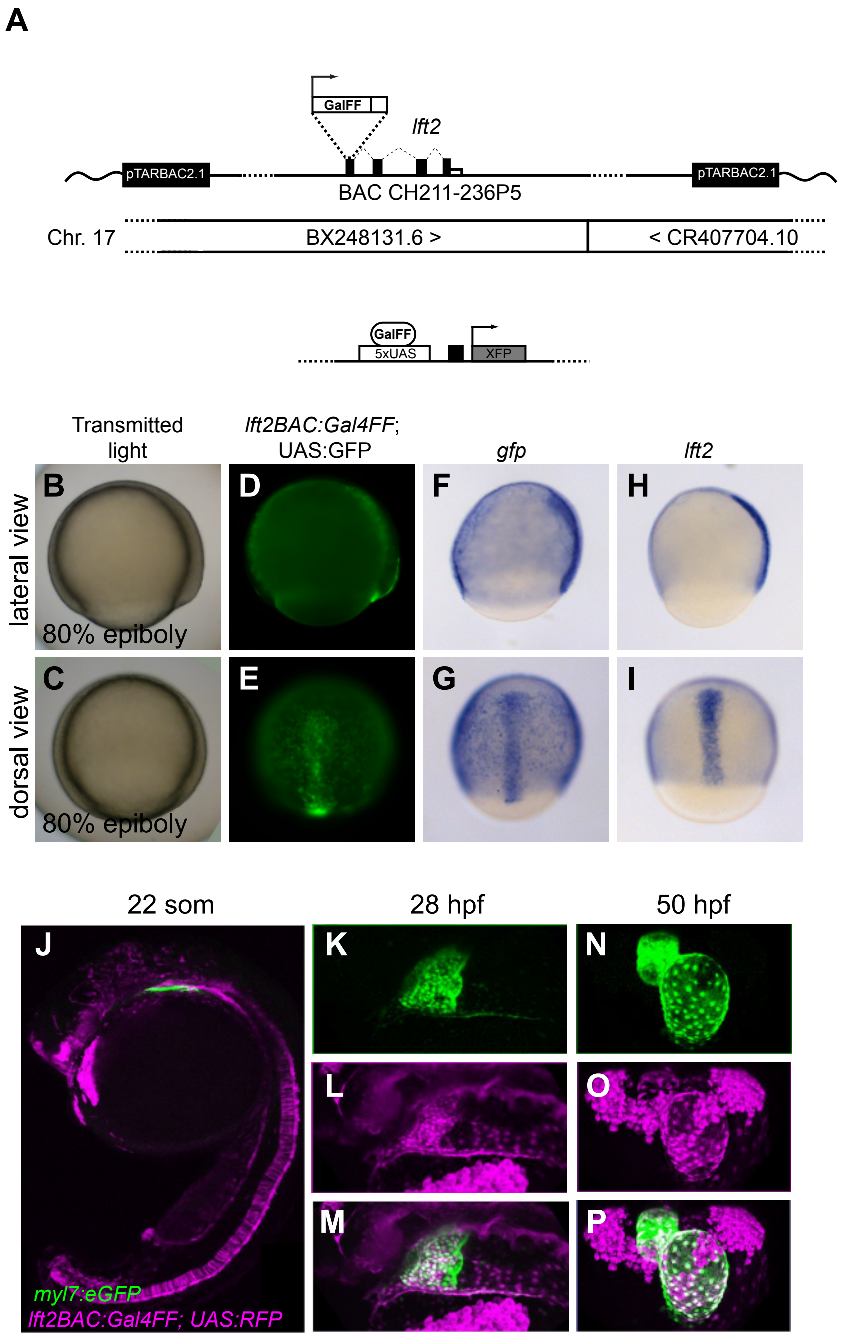

Figure S2 - Cardiac jelly in the atrium is asymmetric at 50hpf

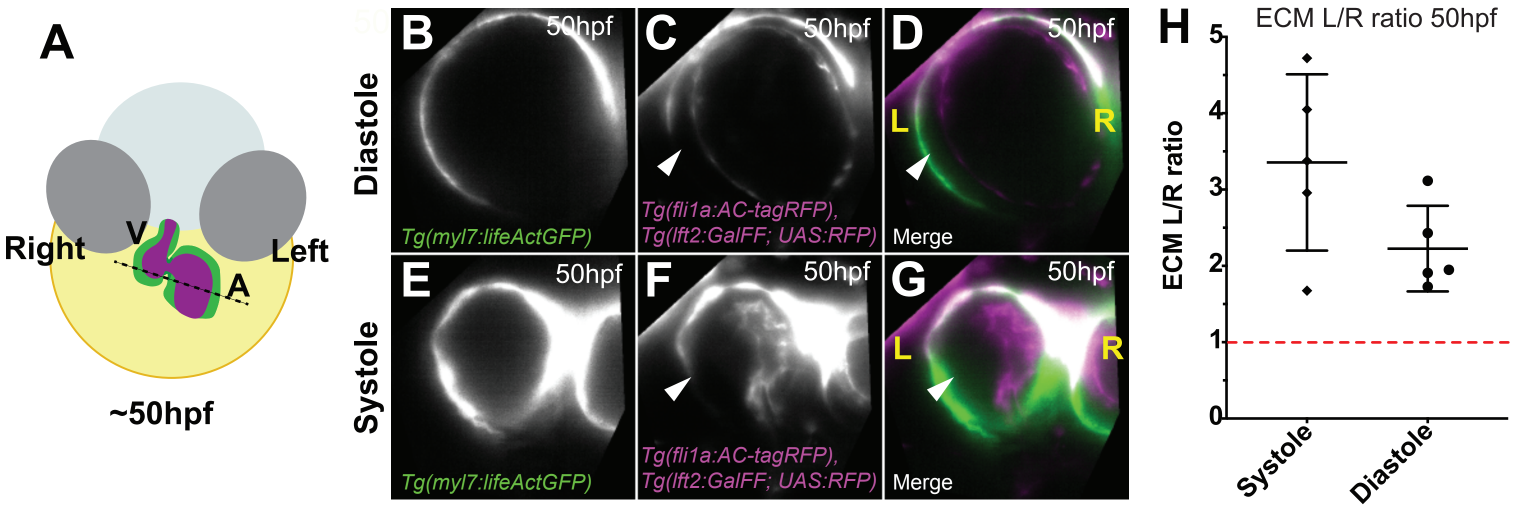

Figure S3 - The ECM constituents HA, *versican* and *aggrecan* are not asymmetrically expressed in the heart tube

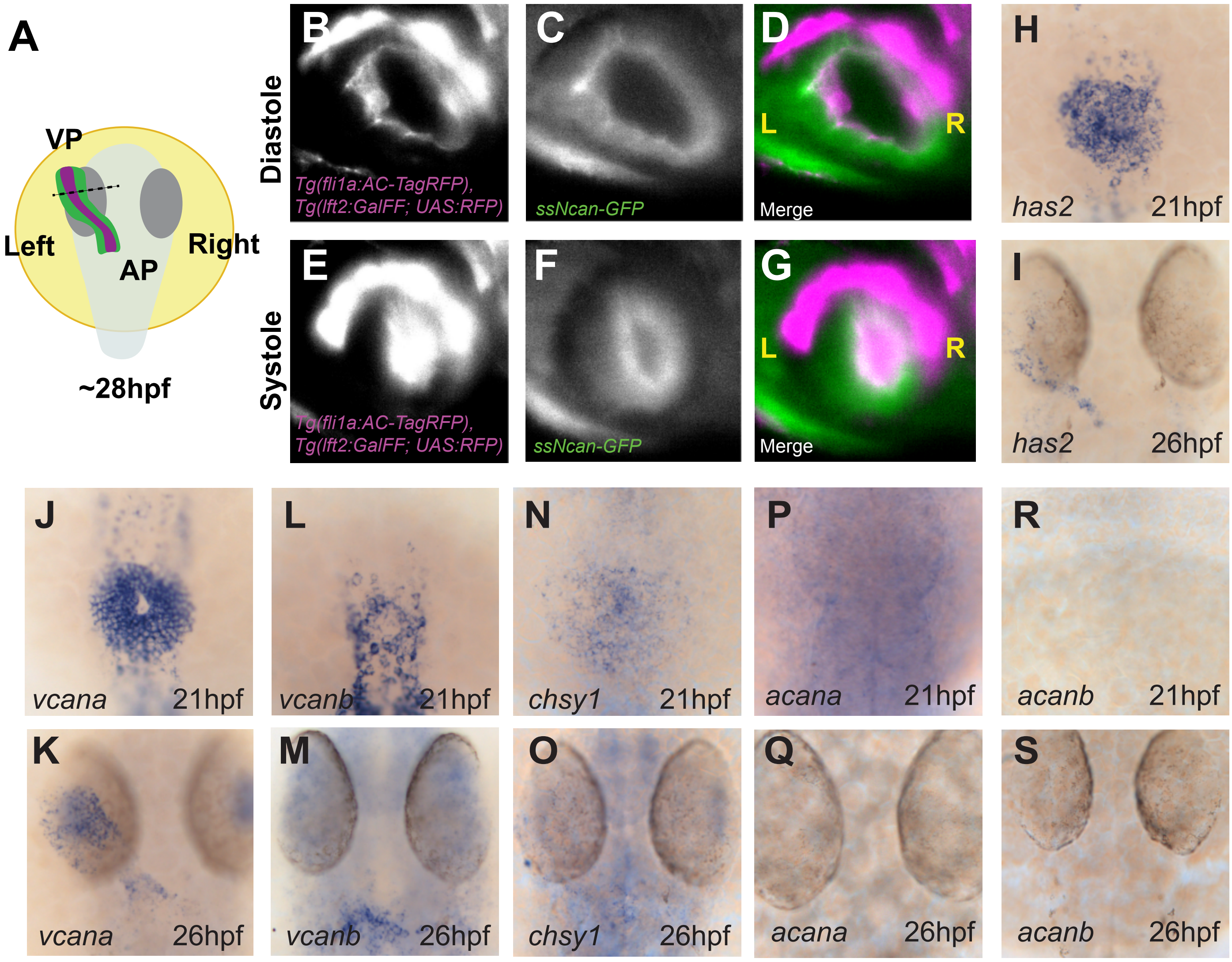

Figure S4 - Hapln1a is required for cardiac morphogenesis

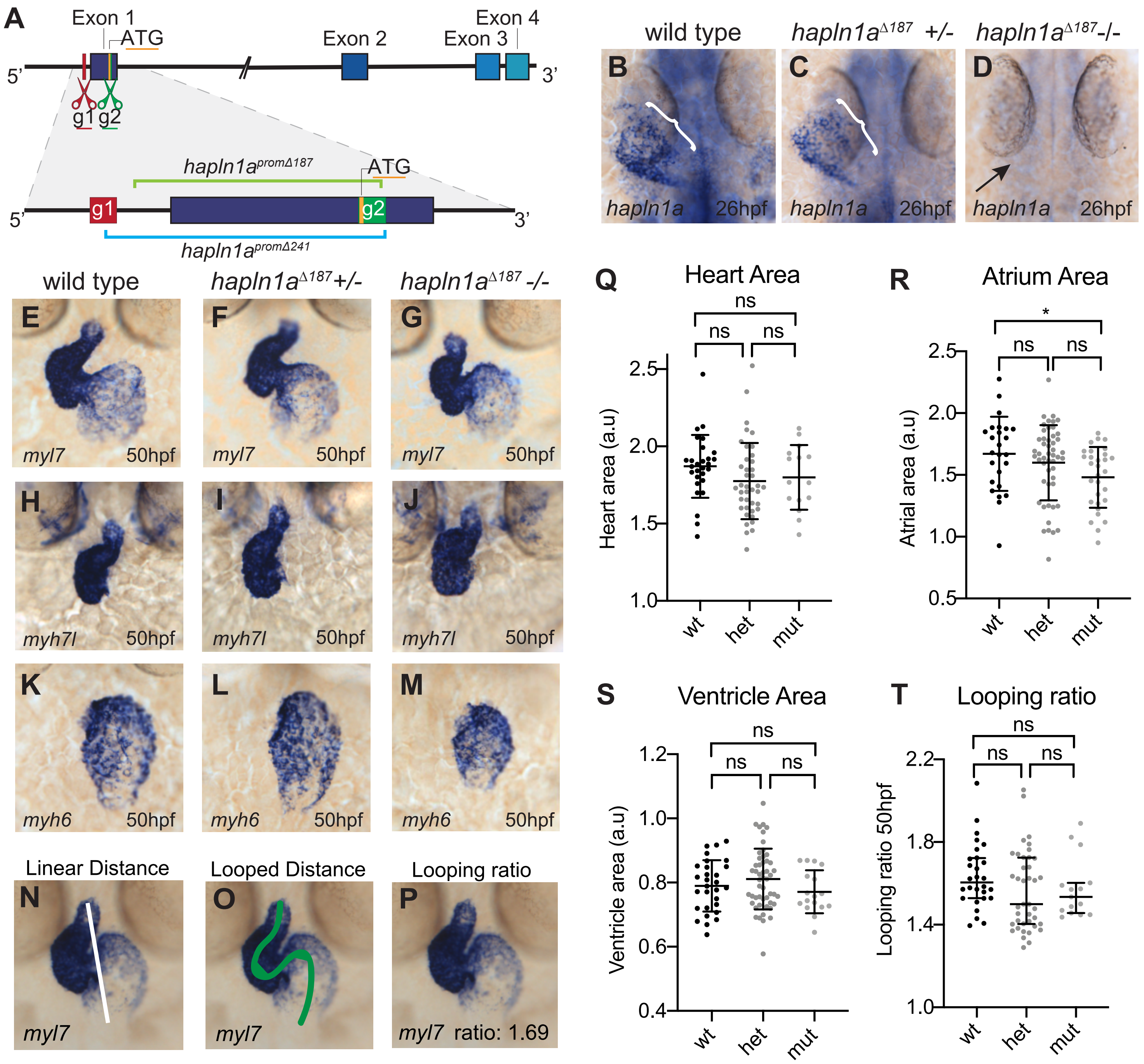

Figure S5 - *hapln1a* morpholino is a loss-of-function model and recapitulates *hapln1a* mutant cardiac phenotypes

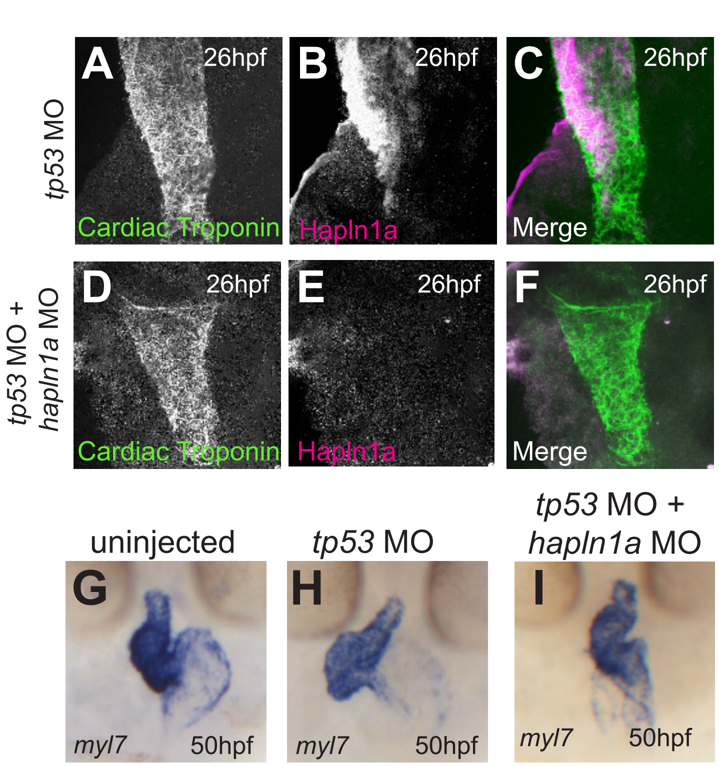

Figure S6 - HA is required at early stages of heart development for cardiac morpho-genesis and interacts with Hapln1a

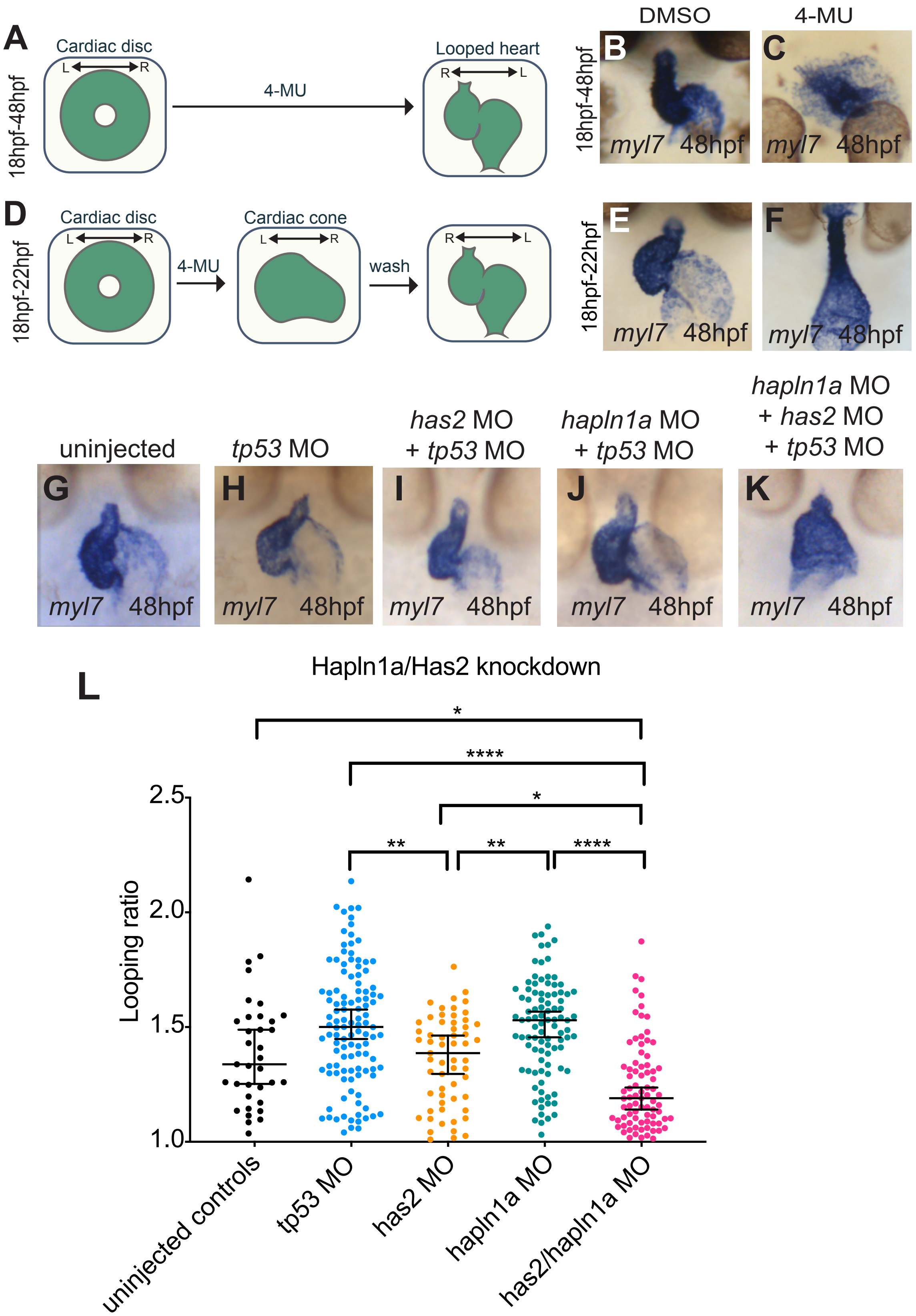
